# Distinct subtypes of astrocytes selectively regulate specific inhibitory synapses

**DOI:** 10.1101/2025.05.16.654545

**Authors:** Darren Clarke, Anthony Bosson, Ève Honoré, Karelle Contant, Elena Avignone, Jean-Claude Lacaille, Richard Robitaille

**Affiliations:** Département de Neurosciences, Faculté de Médecine, Université de Montréal, Montréal, Canada; Groupe de Recherche sur la signalisation neurale et la circuiterie, Université de Montréal, Montréal, Canada; Centre de recherche du Centre hospitalier de l’Université de Montréal, Montréal, Canada; Department of Psychiatry, Douglas Mental Health University Institute, McGill University, Montréal, Canada; University of Bordeaux, INSERM, Neurocentre Magendie, U1215, F-3300 Bordeaux, France; Centre Interdisciplinaire de Recherche sur le Cerveau et l’Apprentissage, Montréal, Canada

**Author notes:** Correspondence should be addressed to: Richard Robitaille, Département de neurosciences, Université de Montréal, PO box 6128, station centre-ville, Montreal, H3C 3J7, Jean-Claude Lacaille, Département de neurosciences, Université de Montréal, PO box 6128, station centre-ville, Montreal, H3C 3J7. denotes equal authorship (Ève Honoré, Karelle Contant). denotes equal senior authorship (Jean-Claude Lacaille, Richard Robitaille).

## Abstract

Astrocyte functional heterogeneity within a given neuronal circuit remains largely undetermined, particularly their role at tripartite synapses. Here, we examine multiple functional characteristics of astrocytes distinguished by their specific spatial relation to inhibitory synapses made on distinct hippocampal CA1 pyramidal cell domains: astrocytes covering the peri-somatic area in *stratum pyramidale* (SP) receiving input from Parvalbumin interneurons, or the apical dendritic area in *stratum radiatum* (SR) innervated by inhibitory inputs from Somatostatin interneurons. Whole-cell dye-filling and confocal imaging of SR astrocytes showed a typical bushy organization of processes while those of SP astrocytes were more polarized, indicating astrocyte morphological heterogeneity. In addition, SP astrocytes formed a smaller, yet polarised syncytium and displayed greater input resistance relative to SR astrocytes. The two populations of astrocyte are functionally different as indicated by their intrinsic Ca^2+^ signaling properties: SP astrocyte Ca^2+^ events had a lower frequency and temporal density, but greater amplitude, relative to SR astrocytes. Using the territorial segregation of inhibitory synapses, we observed that the selective activation using DREADD or blockade with intracellular BAPTA of the two populations of astrocytes regulated inhibitory synapses exclusively in their own syncytial territory. Furthermore, each astrocyte population selectively mediated long-term depression at the respective inhibitory synapses through Ca^2+^-dependent modulation of post-synaptic targets. These results indicate domain-specific regulation of inhibitory synapses by distinct SP and SR astrocyte syncytia. Transcriptional analysis revealed enriched gene expression in SP relative to SR astrocytes, notably for regulation of cell growth and morphology, synaptic function and signaling. Overall, our findings reveal a functional specialization of astrocyte subtypes in the hippocampus, highlighting heterogeneous astrocyte regulation of hippocampal synaptic networks important for learning and memory.

**Graphical abstract:** 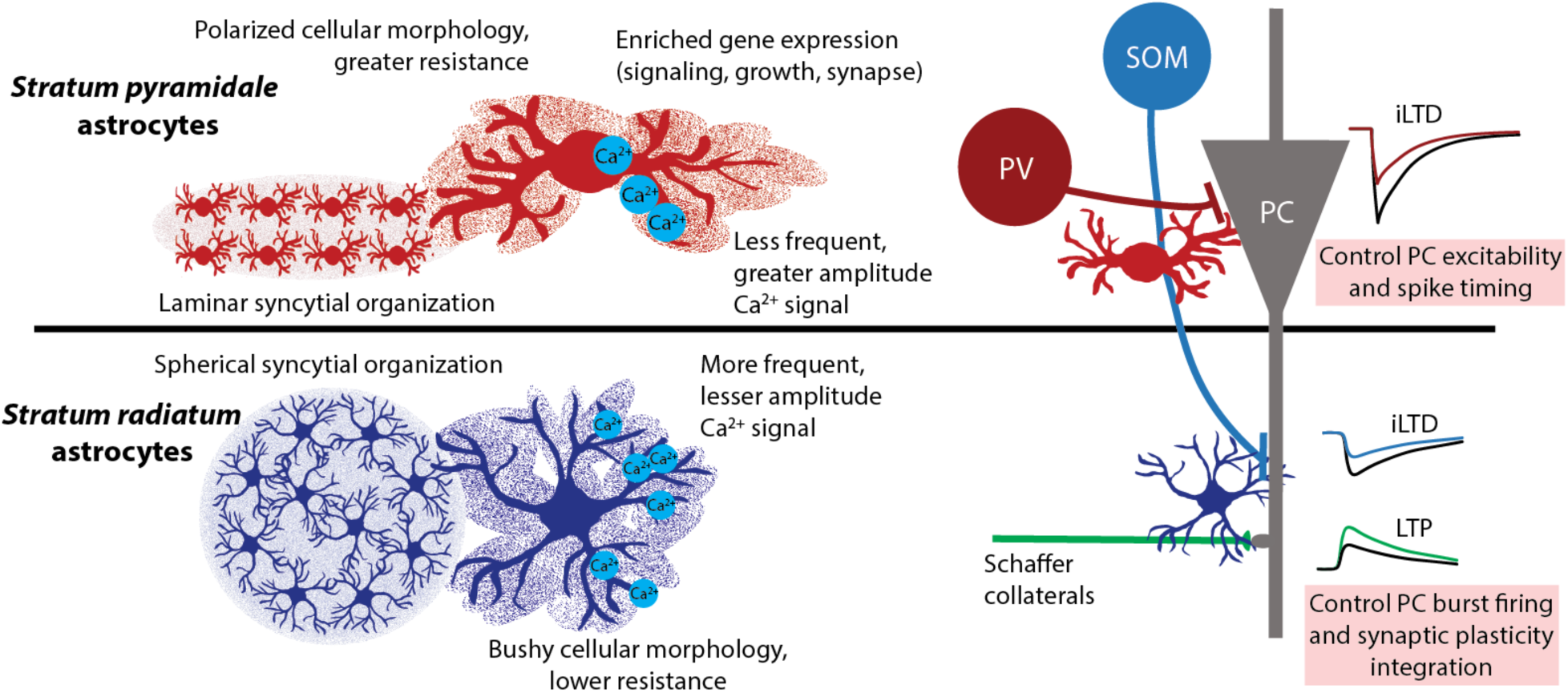

## INTRODUCTION

Astrocytes provide essential support to neurons in the central nervous system through a range of regulatory functions, including modulation of ionic (Orkand et al., 1966, Newman et al., 1984), synaptic (Bergles and Jahr, 1997, Kang et al., 1998, Perea and Araque, 2007), and metabolic activity (Magistretti, 2006). Astrocytes were classically described with heterogenous features (Kimelberg, 2004), such as topographical distribution (Andriezen, 1893) and general physiological properties including morphology, reactivity, GFAP expression (Escartin et al., 2021, Walz and Lang, 1998) but with homologous function. Transcriptional examination of astrocytes has elaborated on molecular heterogeneity of astrocyte populations (Bayraktar et al., 2020, Lanjakornsiripan et al., 2018, Shah et al., 2016). Recent studies highlight the relationship between molecular subtypes, morphological specializations (Endo et al., 2022) and physiological properties between brain regions and within circuits (Batiuk et al., 2020, Chai et al., 2017, Karpf et al., 2022). However, it remains undetermined whether astrocytes *function* heterogeneously within a given neural circuit, and their consequence on the neural circuit. For instance, functional heterogeneity in calcium signaling, a determinant of many astrocyte functions (Henneberger et al., 2010, Kang et al., 1998, Perea and Araque, 2007, Porter and McCarthy, 1996), has been reported for astrocytes of the cerebral cortex in a layer-specific manner (Takata and Hirase, 2008), and cortical astrocytes are distinct from hippocampal astrocytes in both spontaneous and evoked somatic Ca^2+^ signaling dynamics (Batiuk et al., 2020). However, notwithstanding the intertwined relationship between astrocyte Ca^2+^ signaling and synaptic regulation, there is a surprising paucity of evidence of heterogeneity of astrocyte synaptic regulation.

Knowing that astrocytes regulate inhibitory synapses in a Ca^2+^-dependent manner in hippocampus (Deemyad et al., 2018, Kang et al., 1998, Matos et al., 2018, Shen et al., 2022, Shigetomi et al., 2012), we examined the functional heterogeneity of astrocytes by exploiting the laminar organization of the CA1 region of hippocampus, and its territorial inhibitory regulation of pyramidal cells. Amongst the multiple subtypes of inhibitory interneuron that regulate CA1 pyramidal neuron activity (Bezaire and Soltesz, 2013, Cobb et al., 1997, Pelkey et al., 2017, Somogyi and Klausberger, 2005), we exploited two inhibitory cell types owing to their roles and segregated location of their synapses onto pyramidal cells. The Parvalbumin (PV) interneurons synapse peri-somatically to CA1 pyramidal neurons in the stratum pyramidale (SP) to regulate their excitability and spike timing (Cobb et al., 1997, Megı’as et al., 2001, Somogyi et al., 1983). The Somatostatin (SOM) interneurons make synaptic contacts on dendrites of pyramidal neurons in stratum radiatum (SR) and stratum *lacunosum-moleculare* to regulate pyramidal cell burst firing, synaptic plasticity and integration (Halasy et al., 1996, Lacaille et al., 1987, Pouille and Scanziani, 2004, Sik et al., 1995).

In the current study, we assessed physiological and functional properties of astrocytes located in somatic (SP) and dendritic (SR) inhibitory synaptic territories, and their underlying molecular differences. We report that the two astrocyte populations have distinct morphological, syncytial, electrical, and signaling characteristics. Moreover, they selectively and exclusively regulate the transmission and plasticity of a subtype of inhibitory synapse. Accordingly with their putatively different function, SP relative to SR astrocytes have enriched expression of genes involved in regulation of cell growth and morphology, signaling and synaptic function. These results highlight the heterogeneous astrocyte populations of CA1 hippocampus, their specialized functional properties, and their discrete domains of synaptic regulation, suggesting subpopulation-specific roles of astrocytes in regulating hippocampal networks.

## RESULTS

### Astrocyte subtype-specific morphological and biophysical properties

Astrocytes display morphological specializations across brain regions that are informed by the surrounding environment (Cheng et al., 2023, Endo et al., 2022). Hence, we first explored the morphological and network features of astrocytes that occupy different inhibitory synaptic niches in SP and SR of acute slices of dorsal CA1 hippocampus using intracellular labeling with Alexa-Fluor 488 dye (AF488; 150µM) during patch-clamp recordings of astrocytes combined with confocal imaging. While SR astrocytes displayed a typical bushy, spherical morphology, with numerous fine processes (Anders et al., 2014, Viana et al., 2023), SP astrocytes exhibited a polarized appearance, with fewer main processes and less elaborate fine processes (Figure 1A, B). We observed that the polarization (expressed as ratios of astrocyte diameters in parallel with the pyramidal layer (*x*) and perpendicular (*y*) axes) of SP astrocytes was significantly more pronounced than SR astrocytes, along the axis of the pyramidal cell later (Figure 1C, D). Astrocyte polarity shifted from parallel to the pyramidal layer to perpendicular as the distance into SR increased (Figure 1C).

**Figure 1.**
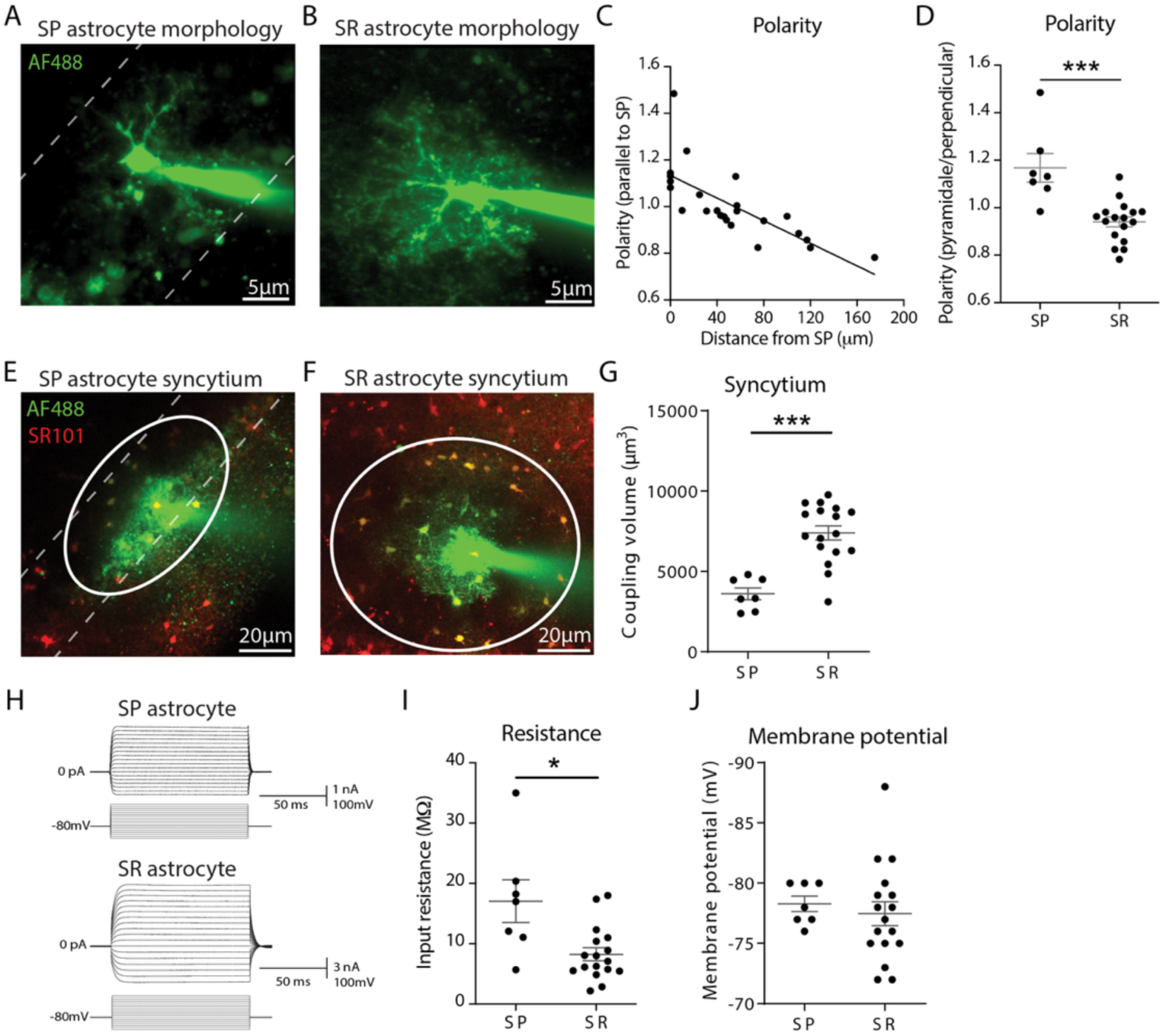
Distinct morphological and biophysical properties of SP and SR astrocytes. A, B, Maximum intensity projection images of fluorescent AF488 dye filled astrocytes illustrates cell morphology of SP astrocytes (A) and SR astrocytes (B). C, D, Graphs of relationship between astrocyte polarity and distance in the SR layer (C, R^2^=0.4459, p<0.001), and of polarity of SP and SR astrocytes (D). E, F, Fluorescence imaging of AF488 and SR101 dye filled astrocyte syncytium for SP astrocytes (E) and SR astrocytes (F), dashed lines indicate SP border. G, Graph depicting syncytium size difference between SP and SR astrocytes. H, Representative example of recordings of astrocyte currents in SP (top) and SR (bottom) astrocytes. I, J, Graphs showing differences in cell input resistance (I) but not resting membrane potential (J) in SP and SR astrocytes (n=7 cells, 7 mice for SP astrocytes; n=17 cells, 8 mice for SR astrocytes). * p<0.05, *** p<0.001, bars represent mean and SEM.

The astrocyte polarity further reflected in their syncytium organization, monitored by AF488 dye diffusing from the patch electrode through gap junctions and colocalization with astrocyte-specific sulforhodamine 101 (SR101, 0.25µM) labeling. SP astrocyte syncytia were spatially restricted along SP with limited extensions up to 20µm into stratum oriens and SR (Figure 1E, G), thus we define SR astrocytes within 20µm of the pyramidal layer as SP astrocytes for these experiments. SR astrocytes displayed typically spherical networks that were significantly larger in comparison and were contained to the SR (Figure 1F, G). Together these findings indicate distinct morphological organization of SP and SR astrocytes at the cell and syncytium levels.

The morphological distinctions of the different astrocytes were further extended in their intrinsic biophysical properties, whereby SR astrocytes had a lower input resistance than SP astrocytes (Figure 1H, I) while maintaining a similar resting membrane potential (Figure 1J).

Together, these results indicate that there are two distinct groups of astrocytes that occupy selective territories in the CA1 hippocampus with distinct intrinsic membrane properties and morphology. Astrocyte subtype-specific Ca^2+^ transients

With the major structural and biophysical properties, we next investigated if the two populations of astrocytes also had different functional properties. We examined whether SP and SR astrocytes had distinct Ca^2+^ signaling dynamics since astrocyte Ca^2+^ signaling is crucial for detection and regulation of neuronal synaptic activity (Aguado et al., 2002, Kang et al., 1998, Panatier et al., 2011, Pasti et al., 1997, Porter and McCarthy, 1996, Nett et al., 2002). Astrocytes were separated in two groups according to their polarity (∼ 50μm away from SP, Figure 1C) and extension of the SP astrocyte syncytium (∼ 20μm away from SP, Figure 1F). SP astrocytes were imaged in *stratum pyramidale* or within 20μm of its border with *stratum radiatum* while SR astrocytes were imaged in *stratum radiatum*, more than 20μm away from the SP border. Representative examples of astrocyte calcium events are shown in Figure 2 (SP astrocyte Figure 2A-C, SR astrocyte Figure 2D-F; Supplementary Video 1).

**Figure 2.**
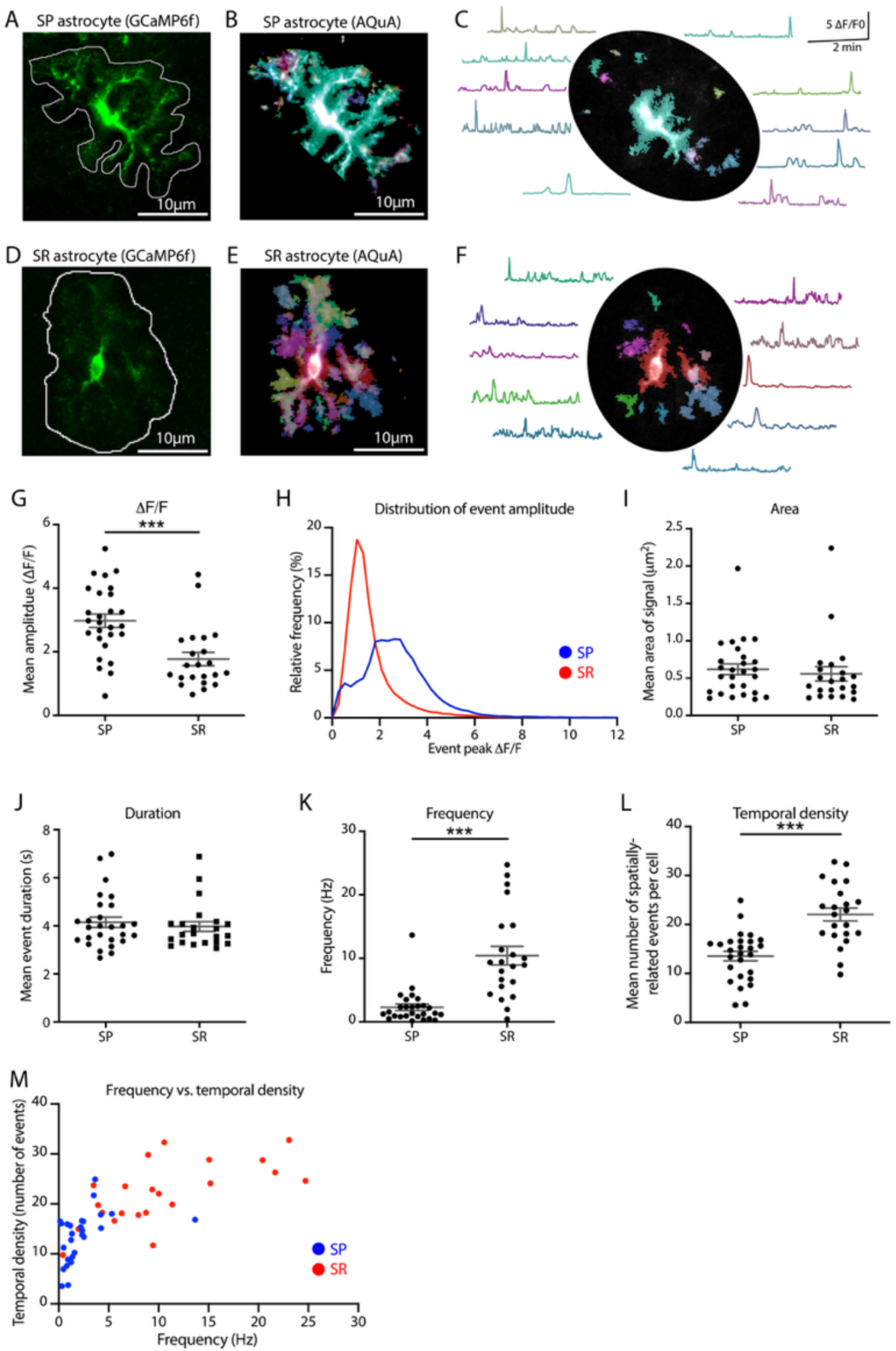
Distinct Ca^2+^ responses in SR and SP astrocytes. A-F, Representative images of Ca^2+^ signaling in SP and SR astrocytes. A, D, Maximum intensity projection images of GCaMP6f expression in SP and SR astrocytes. B, E, Ca^2+^ events detected in AQuA for SP and SR astrocytes. C, F, Example of astrocyte Ca^2+^ transients at selected ROIs for SP and SR astrocytes. See videos (Supplementary Video 1) for visualization of Ca^2+^ transients in astrocytes. G-M, Quantitative analysis of Ca^2+^ activity in SP and SR astrocytes. Peak amplitude (G), relative frequency distribution (H), area (I), duration (J), frequency (K), and temporal density (L) of Ca^2+^ events in SP and SR astrocytes. M, Scatter plot of frequency and temporal density of Ca^2+^ events in SP and SR astrocytes. SP astrocytes n=27 cells, 6 mice; SR astrocytes n=22 cells, 9 mice. *** p<0.001, bars represent mean and SEM.

#### Kinetics

Ca^2+^ events were more frequent in the processes than the soma (Supplementary Video 1), with significantly larger amplitude (ΔF/F) and area under the curve in SP astrocytes than in SR astrocytes (Table 1, Figure 2G, H). Yet, other spatial and temporal features, such as event area (Figure 2I), event duration (Figure 2J), perimeter, rise time and decay time (Table 1), were not significantly different.

**Table 1.**
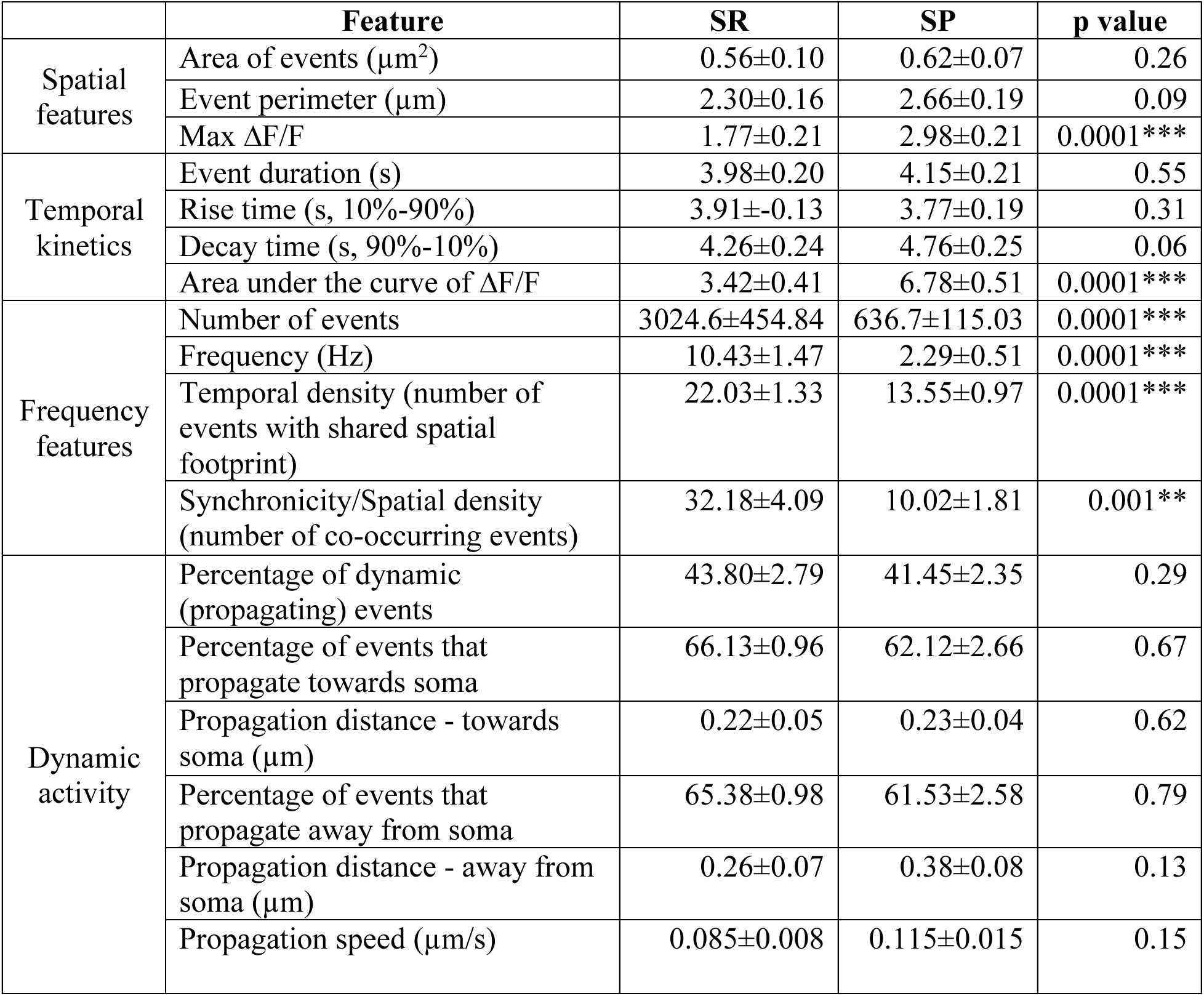
Characteristics and statistical outcome of astrocyte Ca^2+^ events between SR and SP astrocytes. Values listed are mean ± SEM of astrocyte subpopulation acquired from independent recordings of individual astrocytes.

#### Frequency

Frequency of Ca^2+^events, an important factor in astrocyte Ca^2+^ transient properties (Perea and Araque, 2005, Takata and Hirase, 2008), was significantly higher in SR astrocytes than in SP astrocytes (Figure 2K). Furthermore, the temporal density, that is the number of events that share a spatial footprint with another event, was greater in SR astrocytes than SP astrocytes (Figure 2L). A higher temporal density was also observed in SR astrocytes that showed frequency of Ca^2+^ of events like SP astrocytes (Figure 2M).

#### Spatial propagation

Propagation of local Ca^2+^ responses between adjacent compartments contribute to the integration of information by astrocytes (Kang et al., 1998, Kuga et al., 2011, Perea and Araque, 2005). Ca^2+^ events showed limited spatial displacement and, as such, were more frequently static than dynamic for both SP and SR astrocyte populations, with no difference between the two astrocyte groups (Table 1). The direction, mean distance and speed of propagative events were similar between SP and SR astrocytes, with a prevalence for propagating events to occur in the direction towards the soma (Table 1).

Our results indicate that Ca^2+^ signaling differs in SP and SR astrocytes in terms of amplitude, frequency, and spatial relation of the Ca^2+^ events, while the dynamic/propagative features of Ca^2+^ events are similar. Thus, the two populations of astrocytes may be functionally heterogeneous since Ca^2+^ dynamics is a main regulator of astrocyte functions.

### Depression of synaptic inhibition by astrocytes

Considering the spatial differences of the two astrocyte populations, we next assessed their respective modulation of synaptic activity (Araque et al., 2014, Matos et al., 2018, Panatier et al., 2011). We exploited the targeting of specific domains of pyramidal neurons by inhibitory synapses that matched the territories of the two subtypes of astrocytes, that is PV interneurons innervating the perisomatic region covered by SP astrocytes and SOM interneurons innervating the dendritic region covered by SR astrocytes. While SOM interneuron projections can span many layers of CA1 (Pelkey et al., 2017), here we focus on SOM interneuron synapses located in *stratum radiatum*.

PV and SOM inhibitory synapses were selectively activated by optogenetics using Cre-specific expression of Channelrhodopsin2 (EF1a-DIO-hChR2) in PV interneurons (Pvalb-IRES-Cre mice) or SOM interneurons (SOM-IRES-Cre mice), respectively (Figure 3A, B and 3E, F).

**Figure 3.**
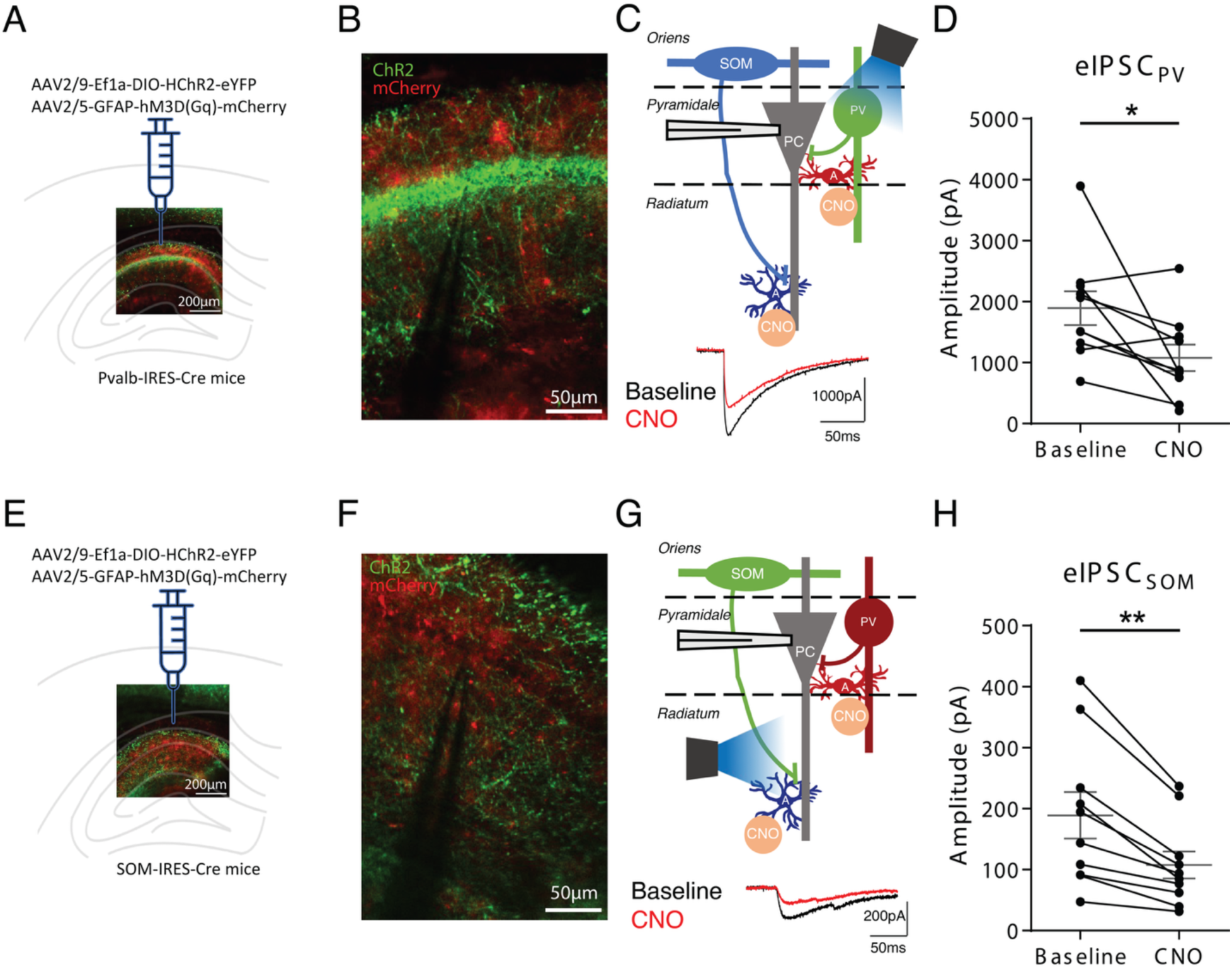
Chemogenetic activation of astrocytes reduces synaptic inhibition of pyramidal neurons evoked by optogenetic stimulation of PV and SOM interneurons. A-C, Fluorescent images (A, B) and diagram of experimental design (C) of expression of hChR2-eYFP in PV interneurons and DREADD-mCherry in astrocytes. D, Summary graph showing depression of eIPSC_PV_ amplitude by CNO (paired t-test, p<0.05, n=10 cells). E-G, Similar experimental design (E, F), and diagram representation (G) for expression of hChR2-eYFP in SOM interneurons and DREADD-mCherry in astrocytes. H, Summary graph showing depression of eIPSC_SOM_ amplitude by CNO (paired t-test, p<0.01, n=10 cells). Dots represent individual cells, mean and SEM presented in grey Representative eIPSC traces are provided in C and G. * p<0.05, ** p<0.01

Prior to testing the relative contribution of each astrocyte sub-types on the regulation of inhibitory synapses, we first determined whether both types of synapses were regulated by astrocytes. Astrocytes were stimulated using G_q_ DREADDs (mCherry-tagged DREADD for G_q_ GPCR (hM3D G_q_)), a robust and reliable approach (Durkee et al., 2019, Nagai et al., 2021). Whole-cell recordings were obtained from CA1 pyramidal cells of acute hippocampal slices to record optogenetically-evoked inhibitory post-synaptic currents (eIPSCs) in the presence of NBQX (10µM) and AP-5 (20µM) (Figure 3C and 3G).

Chemogenetic stimulation of astrocytes by bath perfusion of CNO (10µM, 20 min) reduced the amplitude of eIPSCs from PV interneurons (eIPSC_PV_; 61% of baseline, Figure 3D) and SOM interneurons (eIPSC_SOM_; 58% of baseline, Figure 3H), indicating depression of transmission at PV and SOM interneuron inhibitory synapses onto pyramidal cells by astrocyte activation. Astrocyte depression of eIPSCs was not due to CNO side effects (Manvich et al., 2018) since synaptic inhibition was not depressed in control experiments with expression of mCherry alone (no DREADD) in astrocytes (eIPSC_PV_; 99% of baseline, Supplementary Figure 1). In addition, astrocyte depression of eIPSCs was unlikely caused by virus-induced glial reactivity, as numbers of IBA1-positive cells (Supplementary Figure 2) and GFAP-positive cells (Supplementary Figure 3) were unaltered in injected relative to contralateral control CA1 hippocampus.

### Astrocyte subtype-specific depression of inhibitory synapses

As we observed distinct astrocyte syncytia (Figure 1) that correspond to the respective synaptic territory of SOM and PV interneurons, we next examined whether the two astrocyte populations independently and selectively depress synaptic inhibition determined by their respective syncytial territory. To this end, the intracellular Ca^2+^ activity of the SR or SP astrocyte networks was clamped by patching an astrocyte in whole-cell configuration and introducing the Ca^2+^ chelator BAPTA (20 mM) in the syncytium via the internal solution, prior to optogenetic activation of interneurons.

We first tested if the DREADD-induced astrocyte depression of synaptic inhibition was due to Ca^2+^ elevation in astrocytes. Chelation of Ca^2+^ in SP astrocytes prevented the CNO-induced depression of eIPSC_PV_ (95% of baseline, Figure 4A-C). Similarly, chelation of Ca^2+^ in SR astrocytes (at least 50µm away from SP) blocked the CNO-induced depression of eIPSC_SOM_ (96% of baseline, Figure 4D-F). These results indicate that astrocyte Ca^2+^ activity mediates depression of inhibitory synapses induced by chemogenetic activation of astrocytes.

**Figure 4.**
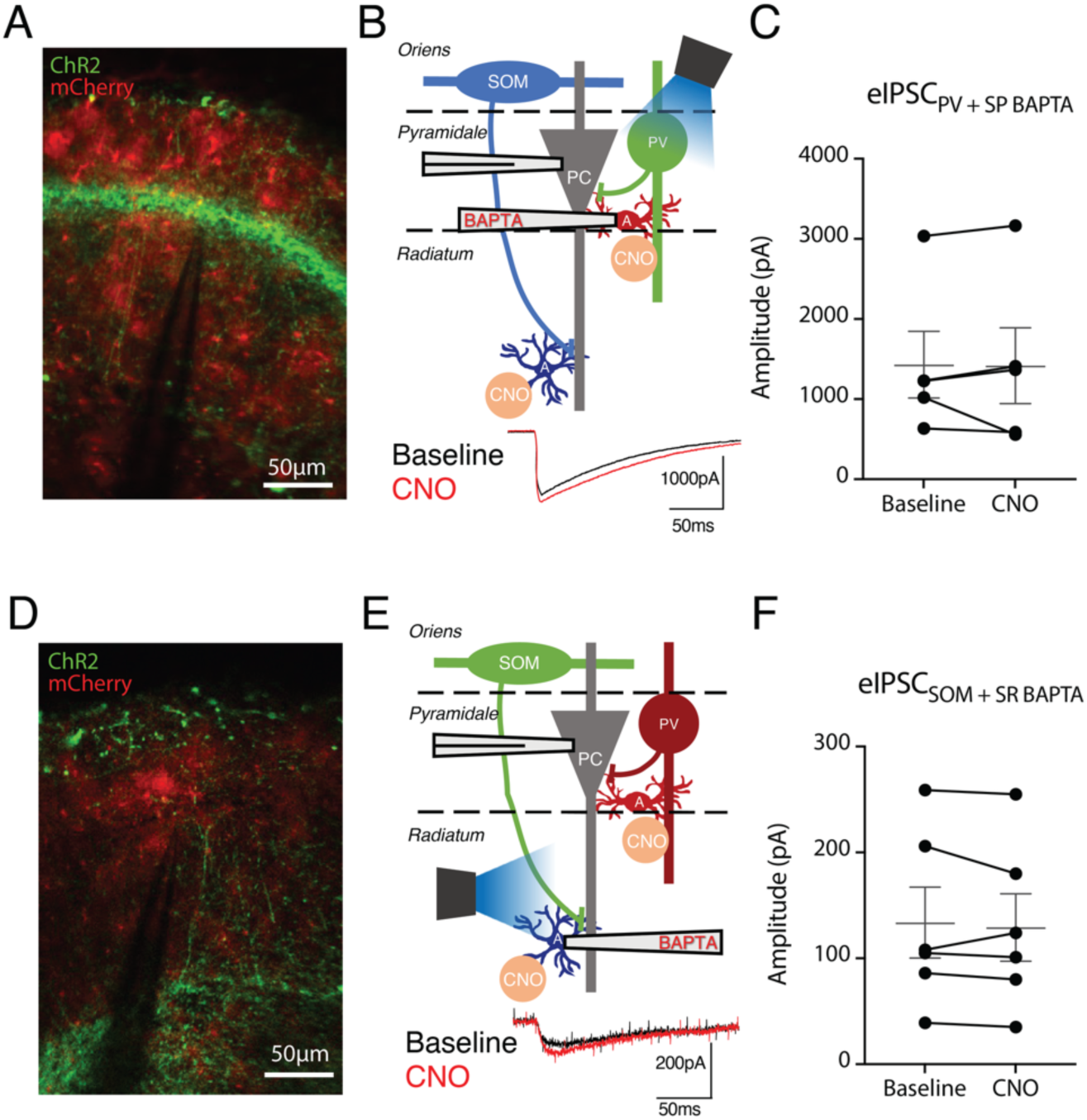
Chelating astrocyte intracellular calcium in the region of activated synapses blocked depression of synaptic inhibition by chemogenetic astrocyte activation. A-C, Fluorescence images (A) and diagram of experimental design (B) of expression of hChR2-eYFP in PV interneurons and DREADD-mCherry in astrocytes, with intracellular BAPTA injection in SP astrocytes. C, Summary graph showing depression of eIPSC_PV_ amplitude by CNO was prevented by BAPTA injection in SP astrocytes (Wilcoxon matched-pairs signed rank, p> 0.05, n=5 cells, 5 mice). D-F, Similar experimental and data representation for expression of hChR2-eYFP in SOM interneurons and DREADD-mCherry in astrocytes (D) with intracellular BAPTA injection in SR astrocytes (E). F, Summary graph showing depression of eIPSC_SOM_ by CNO was blocked by BAPTA injection in SR astrocytes (paired t-test, p> 0.05, n=6 cells, 6 mice). Dots represent individual cells, mean and SEM presented in grey. Representative eIPSC traces are provided in B and E.

Next, we tested whether the astrocytic regulation was confined to their local synaptic area as observed in their exclusive syncytia or extended to distant inhibitory synaptic area. We found that chelation of Ca^2+^ in SR astrocytes did not prevent the depression of PV-evoked eIPSC induced by CNO (47% of baseline, Figure 5A-C). Similarly, chelation of Ca^2+^ in SP astrocytes failed to block CNO-induced depression of SOM-evoked eIPSC (62% of baseline, Figure 5F). These results indicate that astrocyte regulation is confined to their syncytial territory covering local inhibitory synapses and does not cross-over to distant inhibitory synapses. Thus, Ca^2+^-dependent astrocyte syncytium regulation of inhibitory synapses has distinct boundaries, suggesting astrocyte subtype-specific regulation of inhibitory synapses.

**Figure 5.**
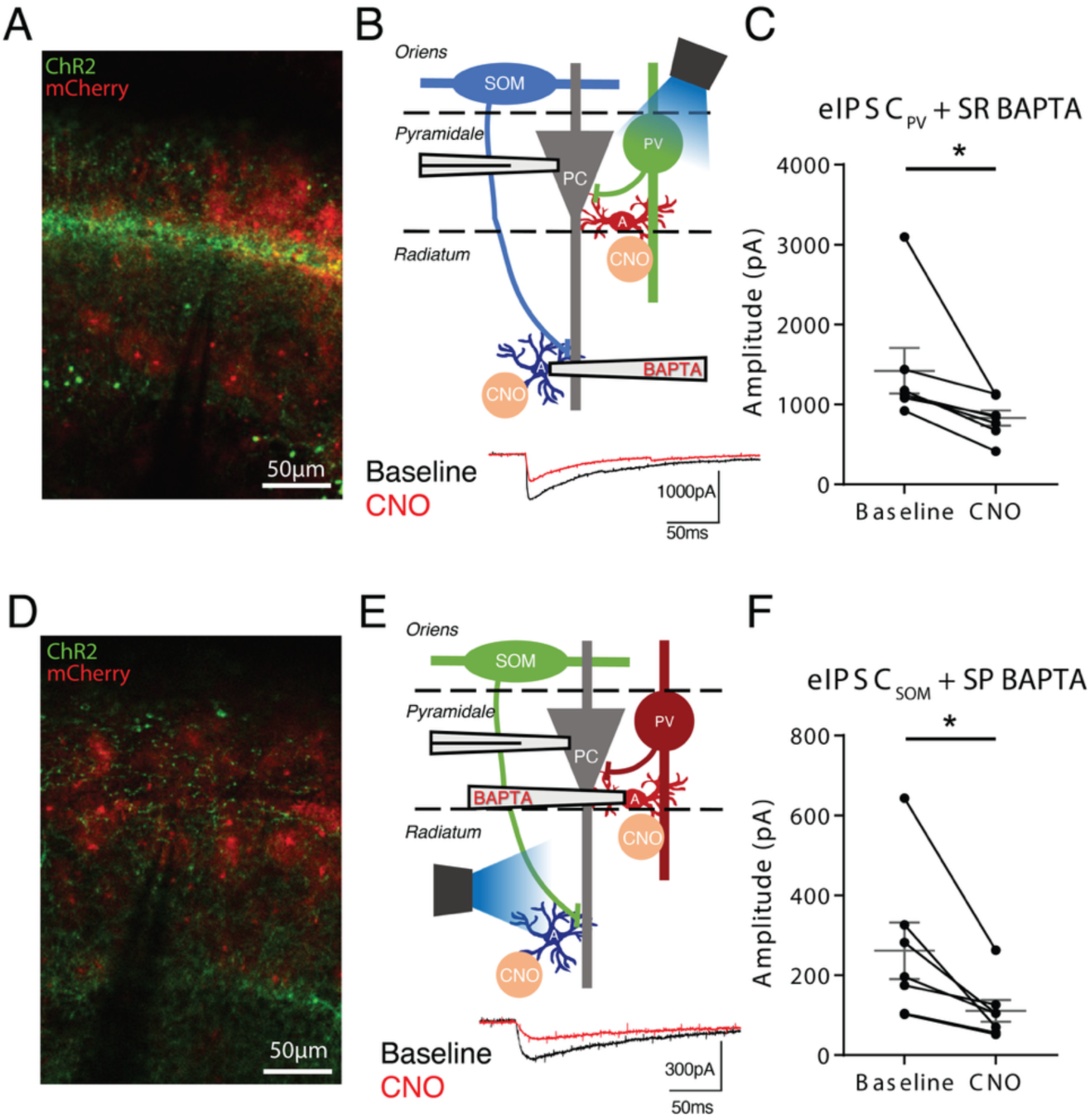
Chelating astrocyte intracellular calcium outside the region of the activated synapses failed to block depression of synaptic inhibition by chemogenetic astrocyte activation. A-C, Fluorescence images (A) and diagram of experimental design (B) of expression of hChR2-eYFP in PV interneurons and DREADD-mCherry in astrocytes, with intracellular BAPTA injection in SR astrocytes. C, Summary graph showing depression of eIPSC_PV_ amplitude by CNO was not affected by BAPTA injection in SR astrocytes (Wilcoxon matched-pairs signed rank, p=0.0156, n=7 cells, 7 mice). D-F, Similar experimental and data representation for expression of hChR2-eYFP in SOM interneurons and DREADD-mCherry in astrocytes (D) with intracellular BAPTA injection in SP astrocytes (E). F, Summary graph showing depression of eIPSC_SOM_ by CNO was not changed by BAPTA injection in SP astrocytes (paired t-test, p=0.021, n=7 cells, 7 mice). Dots represent individual cells, mean and SEM presented in grey. Representative eIPSC traces are provided in B and E. * p<0.05

### Astrocyte subtype-specific long-term depression of inhibitory synapses

The DREADD experiments revealed astrocyte subtype-specific regulation of inhibitory synapses, but they relied on direct and intense chemogenetic stimulation of astrocytes. Hence, we next explored astrocyte regulation of intrinsic activity-dependent long-term plasticity events. We studied activity-dependent plasticity of inhibitory synapses knowing that long-term plasticity at PV and SOM synapses differentially affect pyramidal cell activity and hippocampus function (Royer et al., 2012, Udakis et al., 2020) and that astrocytes are implicated in long-term synaptic plasticity (Henneberger et al., 2010, Kang et al., 1998, Min and Nevian, 2012).

We used the same paradigm as above for optogenetic stimulation of PV and SOM interneurons, and mCherry visualisation of astrocytes while recording eIPSCs in pyramidal cells voltage-clamped at 0mV (Figure 6A, D). For long-term synaptic plasticity, optogenetic theta-burst stimulation (TBS) of PV or SOM interneurons was given and eIPSCs recorded for 30 minutes in the presence of NBQX (10µM) and AP-5 (20µM).

**Figure 6.**
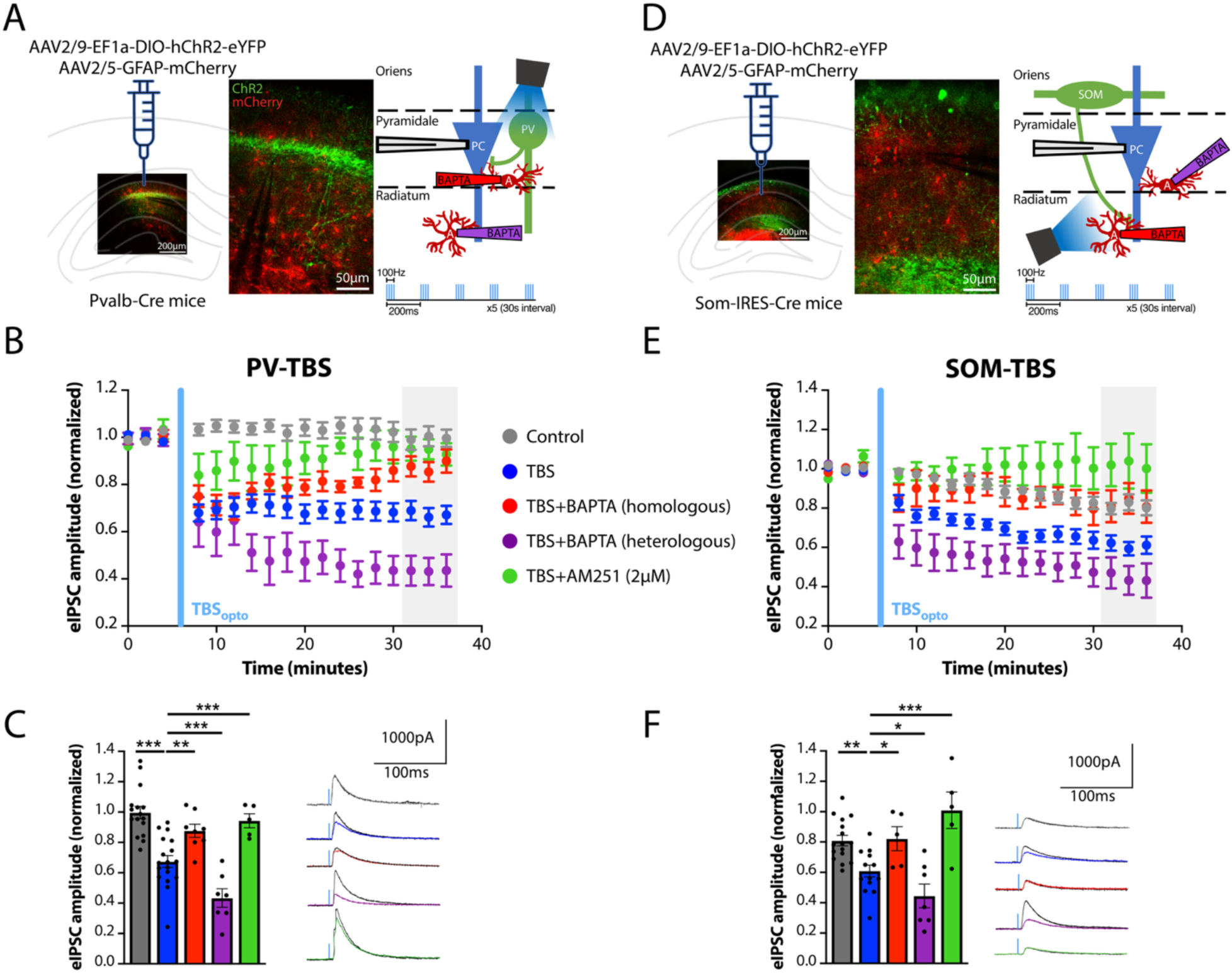
Ca^2+^ signaling in local astrocyte syncytium mediates iLTD at PV and SOM inhibitory synapses. A-C, SP astrocytes mediate iLTD at PV synapses. A, Fluorescent images (left and center panel) and diagram of experimental design (right panel) with expression of hChR2-eYFP in PV interneurons and GFAP-mCherry in astrocytes, intracellular BAPTA injection in SP or SR astrocytes, and optogenetic TBS of PV interneurons. B-C, Summary time plot of eIPSC_PV_ amplitude (B) and graph of eIPSC_PV_ amplitude at 30 min post TBS with representative traces (C), showing no change in responses in control (without TBS, grey, n=16 cells), iLTD induced by optogenetic TBS (blue, n=18 cells, p=0.0001 vs control), block of iLTD by BAPTA injection in SP astrocytes (red, n=8 cells, p=0.0003 vs TBS), no block (but enhancement) of iLTD by BAPTA injection in SR astrocytes (purple, n=7 cells, p=0.0001 vs TBS), and prevention of iLTD by AM251 (green, n=5 cells, p=0.0001 vs TBS). D-F, SR astrocytes mediate iLTD at SOM synapses. D, Similar presentation of images and diagram of experimental design with expression of hChR2-eYFP in SOM interneurons and GFAP-mCherry in astrocytes, intracellular BAPTA injection in SP or SR astrocytes, and optogenetic TBS of SOM interneurons. E-F, Summary time plot of eIPSC_SOM_ amplitude (E) and graph of eIPSC_SOM_ amplitude at 30 min post TBS with representative traces (F), showing run-down of responses in control (without TBS, grey, n=16 cells), iLTD induced by optogenetic TBS (blue, n=14 cells, p=0.0002 vs control), block of iLTD by BAPTA injection in SR astrocytes (red, n=8 cells, p=0.0108 vs TBS), no block (but enhancement) of iLTD by BAPTA injection in SP astrocytes (purple, n=7 cells, p=0.0403 vs TBS), and prevention of iLTD by AM251 (green, n=5 cells, p=0.0001 vs TBS). In B and E, symbols indicate mean ± SEM. In C and F, dots represent individual cells, bars mean ± SEM. Two-way Repeated Measures ANOVA with Bonferroni adjustment for all presented tests. * p<0.05, ** p<0.01, *** p<0.0001

Consistent with previous results (Udakis et al., 2020), TBS induced a long-lasting depression of eIPSC_PV_ (iLTD) relative to pre-TBS baseline (67.4% of baseline; Figure 6B), which was significantly different from control experiments (Figure 6C). eIPSC_PV_ were stable over the 40-minute recording period in control experiments (no TBS stimulation; 99.7% of baseline; Figure 6B). For the SOM interneuron synapses, while a small decrease of eIPSC_SOM_ was observed in control experiments (no TBS, 80.9% of baseline; Figure 6E), TBS induced a long-lasting depression of eIPSC_SOM_ relative to pre-TBS baseline (60.9% of baseline; Figure 6E), which was statistically greater than control experiments (Figure 6F). iLTD of both SOM and PV synapses were of post-synaptic origin since paired-pulse ratio (PPR) was unchanged (Supplementary Figure 4A-B) during iLTD or in control experiments (without TBS). In addition, voltage-clamp of pyramidal neurons at -60mV during TBS prevented iLTD at both PV and SOM synapses, also without changes in paired-pulse ratio (Supplementary Figure 5). Finally, consistent with their postsynaptic origin, iLTD at PV (94.3% of baseline; Figure 6B and C) and SOM synapses (100.9% of baseline, Figure 6E and F) was blocked by the CB1R antagonist AM251 (Araque et al., 2017, Chevaleyre and Castillo, 2003, Chevaleyre et al., 2006). Thus, iLTD at PV and SOM synapses appear of post-synaptic origin and to involve endocannabinoid signaling.

We next tested the astrocyte contribution in iLTD using intracellular Ca^2+^ buffering with BAPTA injection in astrocytes of local and distant synaptic regions. For PV interneuron synaptic plasticity, iLTD of eIPSC_PV_ was blocked by BAPTA injection in SP astrocytes (87.7% of baseline; Figure 6B-C). Importantly, consistent with the territorial astrocyte selectivity, intracellular BAPTA injection in distant SR astrocytes did not block iLTD of eIPSC_PV_ (43.4% of baseline; Figure 6B-C). In accordance with a post-synaptic nature of iLTD, paired-pulse ratio was unchanged in these experiments (Supplementary Figure 4). These results indicate that iLTD induced by TBS at PV inhibitory synapses is mediated by Ca^2+^ signaling in the local SP astrocyte syncytium.

Analogously for SOM interneuron synaptic plasticity, iLTD of eIPSC_SOM_ was blocked by BAPTA injection in SR astrocytes (84.5% of baseline; Figure 6E-F) but not by intracellular BAPTA injection in distant SP astrocytes (44.5% of baseline; Figure 6E-F). Also consistent with a post-synaptic nature of iLTD, paired-pulse ratio was unchanged in these experiments (Supplementary Figure 4). These data indicate that iLTD induced by TBS at SOM inhibitory synapses is mediated by Ca^2+^ signaling in the local SR astrocyte syncytium. Overall, our results reveal a territorial and selective mediation of long-term plasticity of inhibitory synapses by two populations of functionally different astrocytes.

### Astrocyte subtype-specific gene expression

As molecular heterogeneity of hippocampal astrocytes has been suggested (Batiuk et al., 2020, Karpf et al., 2022), we next used transcriptomic analysis to examine differences in gene expression in the astrocyte populations. We probed the publicly available high-resolution spatial Allen Brain Cell atlas of whole mouse brain obtained using MERFISH (Yao et al., 2023) to reconstruct SP and SR astrocytes of dorsal CA1 (shown in two- and three-dimensional representations respectively, Figure 7A, Figure 7B). However, the spatial distribution of SP astrocytes in the Allen Brain Cell atlas is restricted to their conventional categorization based solely within the pyramidal layer. Hence, based on our functional data, we expanded SP definition by adding a 20 μm zone that we separately classified as Peri-SP astrocytes (n=199 SP cells, 194 Peri-SP cells, 1514 SR cells). First, the expression of 500 genes for each astrocyte was visualized using a Uniform Manifold Approximation Projection (UMAP, Figure 7C), demonstrating astrocytes discernibly clustering within their layer groups. A heatmap was generated to confirm enrichment of canonical astrocyte markers and low expression of markers for other cell types (Figure 7D, cell type markers enlarged, all 500 genes shown). The cells displayed high expression of *Gfap*, *Gjai*, *Aqp4*, *Sox2* and other genes commonly enriched in astrocytes, and low expression of genes typically expressed in other glial cells and neurons, including *Calb2*, *Reln*, *Th*, and *Opalin*.

**Figure 7.**
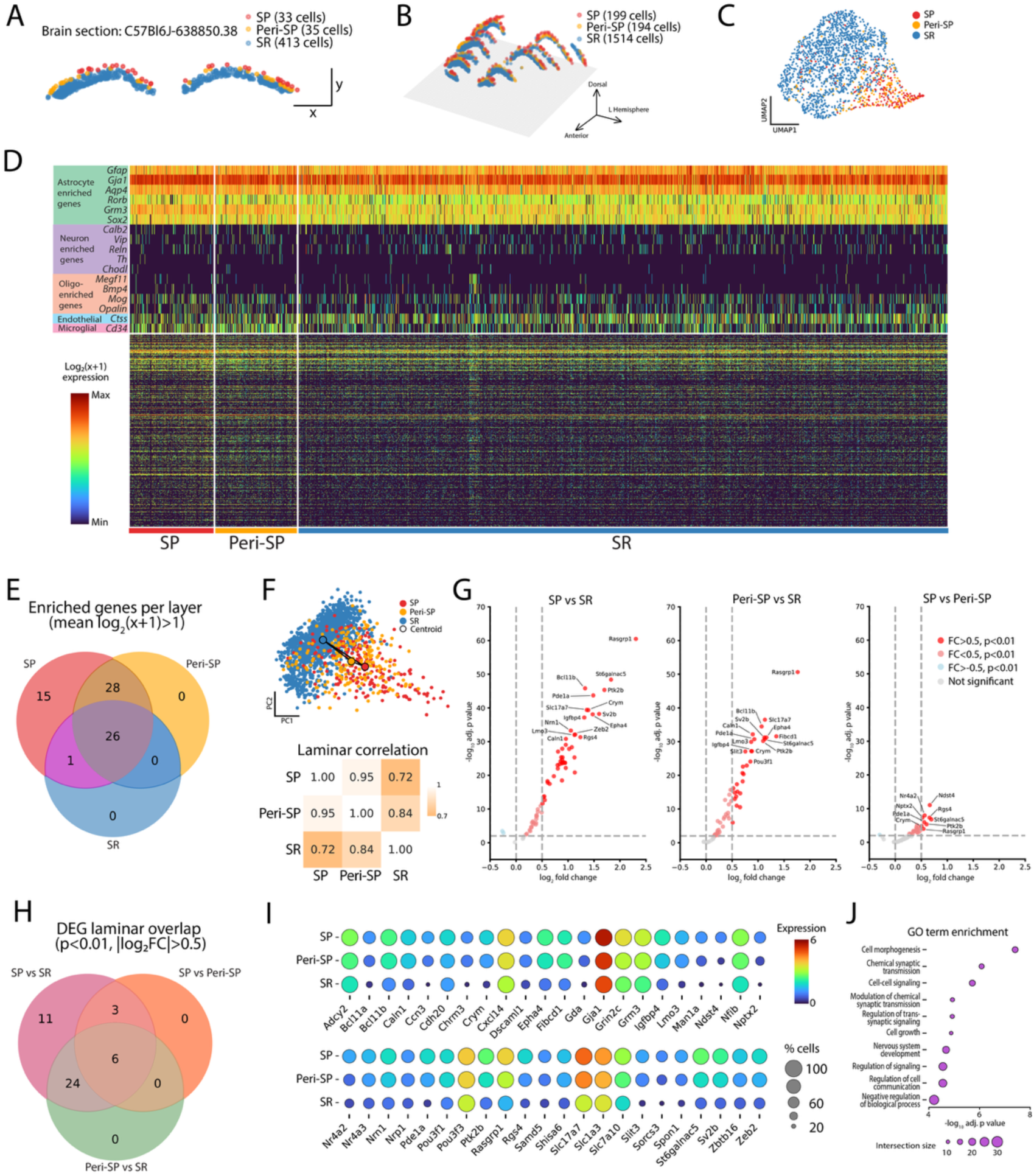
Differential gene expression in SP and SR astrocytes. Transcriptomic analysis of cells in the MERFISH dataset of the Allen Brain Cell Atlas. Reconstruction of probed astrocytes distribution in two- (A) and three-dimensional (B) space, with astrocytes identified as SP, SR, or Peri-SP (SR astrocytes within 20µm of SP). C, SP, Peri-SP, and SR astrocytes were visualized based on gene expression to illustrate heterogeneity between astrocyte populations. D, Gene expression heatmap for 500 genes tested in all cells, with magnified emphasis on select cell-type specific markers to confirm the molecular identity of the cells as astrocytes and not other brain cell types. Rows correspond to genes, columns to cells. E, Diagram showing distribution of 70 enriched genes across astrocyte groups. F, Assessment of expression similarity across astrocyte groups. Top, principal component analysis (PCA) from 70 enriched genes, each dot represents a cell, group centroids depicted by black circle. Bottom, plot of Spearman correlation of mean expression for each enriched gene between astrocyte populations, lighter color represents stronger correlation. G, Volcano plots comparing gene expression between astrocyte groups. Threshold requirement set at p<0.01, |log_2_(x+1) fold change| > 0.5. Top 15 genes with strongest change in expression are identified. H, Summary diagram displaying number of differentially expressed genes (DEGs) across astrocyte groups, illustrating the SP and Peri-SP overlap in DEG relative to SR. I, Dot plot illustrating gene expression level (encoded by circle color) and percentage of cells with transcript expression (encoded by circle diameter). J, Dot plot of ten gene ontology (GO) terms obtained by significant genes considered in SP vs SR, ranked by p value.

We next considered enriched gene expression similarities and differences between astrocyte populations. Enriched expression was observed for 70 genes in SP astrocytes, 54 genes in Peri-SP astrocytes, and 27 genes in SR astrocytes (Figure 7E). The 70 SP genes encompassed all genes enriched in Peri-SP and SR (77% and 37% overlay with SP respectively), there were no enriched genes unique to Peri-SP or SR. We assessed similarities in enriched gene expression between astrocyte populations by measuring the distance between centroids from principal component analysis (PCA), and Spearman correlation of gene expression vectors. The Euclidean distance in the PCA between the center of SR astrocytes and the center of SP (7.32, Figure 7F) top) and with the center of Peri-SP centers (5.10) was greater than the one between SP and Peri-SP centers (2.25). In addition, the Spearman correlation demonstrated a closer association between SP and Peri-SP than SR to either population (Figure 7F, bottom). Together these findings indicate molecular similarity between SP and Peri-SP astrocytes, but transcriptional heterogeneity between them and SR astrocytes.

We next examined differentially expressed genes (DEGs) between astrocyte populations. Forty-four DEGs displayed a significantly greater fold change in SP astrocytes relative to SR astrocytes (Figure 7G; left panel, top 15 genes annotated), including the gap junction protein gene *Gja1,* the adenylate cyclase family gene *Adcy2*, and cell adhesion-related genes *Cdh20* and *Dscaml1*. Furthermore, relative to SR astrocytes, both SP and Peri-SP have enriched expression of receptor and transporter genes *Chrm3*, *Grin2c*, Grm3, *Shisa6*, *Slc17a7*, *Slc1a3* and *Slc7a10*, and a range of development- and plasticity-related genes such as *Bcl11a*, *Epha4*, *Nrn1*, *Slit3*, *Spon1*, and *Zeb2* (Figure 7G, left and middle panels). Consistent with their molecular similarity, only nine DEG were found between SP and Peri-SP astrocytes, six of which were present in every comparison (*Crym*, *Pde1a*, *Ptk2b*, *Rasgrp1*, *Rgs4*, *St6galnac5*; Figure 7G, right panel). In addition, Peri-SP and SR astrocytes shared 30/44 DEG established in the SP-SR comparison, with no DEG unique between Peri-SP and SR (Figure 7H). The gene expression proportions and mean for the 44 DEG is presented in Figure 7I and provided in Supplementary Table 2. Overall, the DEG analyses reveal transcriptional heterogeneity between SP and SR astrocytes. Lastly, we utilized gene ontology to explore the functional relevance of astrocyte subtype-specific differential gene expression. The top terms were related to cell morphogenesis, signaling and synaptic transmission, development, and the regulation of signaling (Figure 7J, Supplementary Table 3). Thus, based on SP and SR astrocyte functional heterogeneity, we unraveled precise molecular specializations that may underly hippocampal astrocyte heterogeneity and their neuronal circuit regulation.

## Discussion

We utilized the territorial segregation of different inhibitory inputs to CA1 pyramidal neurons to explore functional properties of astrocyte heterogeneity. We posited that different synaptic environments would drive physiological and functional distinction in astrocyte populations. We found that CA1 SP and SR astrocytes have different physiological and functional properties, and the discrete syncytia of SP and SR astrocytes exclusively exert control of the inhibitory synaptic territory in their immediate network environment. Furthermore, the functional heterogeneity unraveled an underlying molecular identity of each astrocyte population as determined by their specific gene expression profiles. These findings highlight the compartmentalization and specialization of astrocyte populations within central neuronal circuitry to regulate and support neuron function.

### Functional territorial glial boundaries

Astrocyte populations have been traditionally classified by their location and morphology in nervous system tissue, changes during pathology, and molecular profiles (Khakh and Deneen, 2019, Kimelberg, 2004, Zhang and Barres, 2010). Here we observed a precise organization of astrocytes based on multiple physiological and functional properties. This allowed us to define two distinct functional populations of astrocytes and delineate their location boundaries in CA1. Importantly, these functional distinct identities were further confirmed by our transcriptomic analyses of SP and SR astrocytes that revealed different molecular identities aligned with the physiological specializations of astrocytes found within PV and SOM synaptic environments. Our physiological experiments allowed us to uncover a 20µm SP/SR layer boundary, ultimately instructing our approach to examine and redefine the molecular heterogeneity in CA1 astrocytes. This territorial segregation permits the SP astrocytes to control the peri-somatic environment of pyramidal neurons, whereas the syncytia of the SR astrocytes regulate their dendritic compartment. Thus, our data suggest that astrocytes form exclusive glial enclaves surrounding pyramidal cells: the pyramidale astrocytes creating a unique environment around the peri-somatic area that display a polarized cell and syncytium structure; leaving the dendritic area under the control of the radiatum astrocytes that show a more typical bushy structure and syncytial organization. These exclusive syncytia imply the presence of selective and tailored glial regulation of exclusive synaptic domains that likely instruct the local Ca^2+^ dependent regulation of inhibitory synapse activity we observed following selective BAPTA blockade in DREADD and iLTD experiments. The importance of astrocyte transcriptional regulation to their morphological specialization and functional consequences has previously been established across different brain regions (Endo et al., 2022). Here, we find that molecular heterogeneity of astrocytes within a brain region is consistent with different functional properties of these cells, including structural elements pertaining to their syncytial organization (*Gjai*), and regulation of Ca^2+^ signaling (*Adcy2*, *Caln1*, *Chrm3*, *Fibcd1*). Hence, we argue that the interaction of inhibitory synapses and astrocyte during maturation (Cheng et al., 2023) may instruct astrocytes to specialize their regulatory functions to suit distinct synaptic environments and modulate more proficiently the CA1 circuit.

### Astrocyte heterogeneity beyond their molecular signature

Our work shows that astrocyte heterogeneity goes beyond their molecular signature as we found multiple physiological and functional distinctions between SP and SR astrocytes. In the CA1 hippocampus, there are precise inputs to pyramidal neurons, where an almost exclusive inhibitory innervation arrives peri-somatically in the SP, while dendrites in the SR receive rich and diverse synaptic contacts including excitatory and inhibitory inputs (Cobb et al., 1997, Megı’as et al., 2001). Hence, the difference in synaptic environments could drive the functional differences of the SP and SR astrocytes whereby the rich diversity of excitatory and inhibitory inputs in the SR may promote increased Ca^2+^ activity while the spatial constraints of pyramidal neuron somata and the postnatally developed perineuronal nets may constrain the morphology of SP astrocytes and their additional specialization towards the regulation of inhibitory synapses. While SP and SR astrocytes occupy discrete spatial territories, their physiological and molecular differences are graded within continuous variables. Therefore, in defining functional populations of astrocytes, multiple intrinsic features from molecular to function must be considered in addition to extrinsic influences. This integration of physiological, functional, and molecular approaches to assess heterogeneity would inform whether astrocyte subtypes are discrete or continuous “phenotypes”, and how astrocyte heterogeneity befits function.

Multiple intrinsic and extrinsic factors may contribute to this detailed level of heterogeneity. For instance, astrocyte gene expression influences their morphology (Endo et al., 2022), indicating an intrinsic driver of astrocytic diversity. In addition, neuronal activity and milieu also shape astrocyte morphology (Cheng et al., 2023, Farmer et al., 2016, Tewari et al., 2024), suggesting that the synaptic environment tailors astrocyte specialization (Endo et al., 2022). Indeed, preferential proximity between astrocytes and somata of specific interneuron subtypes has been previously reported in hippocampus and striatum, with implications for regulatory functions (Refaeli et al., 2021, Stedehouder et al., 2024, Yamazaki et al., 2005).

Our findings establish novel levels of functional heterogeneity for astrocyte populations and highlight the importance of contextual classifications such as synaptic environments over anatomical positioning. Indeed, tailored functional astrocyte heterogeneity can be such that it goes beyond their molecular differences at the circuit level that are relative and not absolute, rendering the precise targeting of these specific populations unattainable by current molecular tools. In this manner, we posit that a new frontier has been reached in the study of astrocyte heterogeneity. The currently available physiological, pharmacological, imaging, and molecular tools require precision beyond presently practicable to parse or exploit astrocyte functional heterogeneity to further appreciate their functional significance.

### Astrocytes as adapted synaptic integrators

The excitatory innervation of CA1 is rich in density (Gulyás et al., 1999, Megı’as et al., 2001) and functional significance (Brun et al., 2002, O’Keefe and Dostrovsky, 1971, Wilson and McNaughton, 1993). Previous studies have shown the importance of *stratum radiatum* astrocytes in regulating the transmission and plasticity at CA1 excitatory synapses (Henneberger et al., 2010, Navarrete et al., 2012, Panatier et al., 2011, Perea and Araque, 2007), and their involvement in hippocampal behavior (Doron et al., 2022). However, the pyramidale astrocytes are interacting almost exclusively with PV inhibitory synapses. Hence, this implies that the synaptic integration of the two types of astrocytes is fundamentally different, with the radiatum astrocytes handling highly diversified and extensive synaptic connectivity, while pyramidale astrocytes need to compose almost exclusively with inhibitory synapses. This major difference in the integrative functions of the two types of astrocytes may underlie their different Ca^2+^ activity illustrated by the larger Ca^2+^ responses in pyramidale astrocytes while, albeit smaller, Ca^2+^ responses are more frequent and spatially dynamic in radiatum astrocytes. The distinct integrative functions may also be influenced by differential contribution of channels and receptors (such as GAT3 and GABA_B_R) towards astrocyte regulation of inhibitory synapses (Matos et al., 2018). The architecture of astrocyte compartments is an important factor in Ca^2+^ signal propagation through astrocytes (Lines et al., 2023), and the distinct morphologies of SP and SR astrocytes may structure the different Ca^2+^ signaling profiles and electrical properties we observed. Finally, striatal and hippocampal astrocytes display distinct Ca^2+^ signaling dynamics (Chai et al., 2017), suggesting a role in the function of the brain circuit in shaping astrocyte physiology. Therefore, functional specialization of astrocytes appears precisely adapted to their specific environments, positioning astrocyte populations to regulate their local environment in a manner befitting the requirements of synaptic activity and demands of the circuitry.

### Distinct astrocytes with common neuronal targeted mechanisms

Our work unraveled the astrocytic territorial regulation of spatially distinct inhibitory synapses. In addition, we also showed that astrocytes exclusively mediated the synapse-specific iLTD of their respective synaptic territory through post-synaptic mechanisms (PPR unchanged, depolarization- and CB1R-dependent). This implies that SP and SR astrocytes, through Ca^2+^ dependent pathways, independently modulate post-synaptic elements of inhibitory synapses within the local astrocyte network to produce and maintain iLTD through interactions with endocannabinoid signaling. This suggests that, despite the diversity in astrocyte functional properties such as syncytial organization or Ca^2+^ excitability, their functional output appears to influence the same mechanisms at post-synaptic sites on the neuron. Hence, the strategy to support such a precise functional heterogeneity may be not to alter the fundamental functions of astrocytes but to rely on similar mechanisms to regulate their respective, exclusive domains of inhibitory synapses.

### Impacts of astrocytic heterogeneity on the regulation of hippocampal inhibitory circuits

Astrocyte regulation of CA1 inhibitory synapses has been previously reported at both synaptic and network levels. For example, astrocytes potentiate inhibitory transmission from SR interneurons via Ca^2+^-dependent glutamatergic release (Kang et al., 1998) and drive barrage firing of CA1 neuropeptide-Y interneurons (Deemyad et al., 2018). Furthermore, SR astrocytes potentiate SOM inhibition of CA1 pyramidal neurons via Ca^2+^-dependent ATP release (Matos et al., 2018, Shen et al., 2022), but not PV inhibition. In addition, optogenetic activation of CA1 astrocytes increased CCK interneuron (but not PV) firing frequency and altered *ex vivo* hippocampal oscillations (Tan et al., 2017). Together, this highlights the potent regulatory functions of astrocytes on select inhibitory CA1 circuitry.

As CA1 interneurons regulate synaptic integration and excitability of pyramidal neurons, functional heterogeneity may position astrocyte populations to selectively govern specific properties of the neuronal circuits (Murphy-Royal et al., 2023), thus optimizing learning and memory function of the hippocampus. For instance, through their regulation of PV synapses, pyramidale astrocytes could influence the excitability and spike timing of CA1 pyramidal neurons (Cobb et al., 1997, Megı’as et al., 2001, Somogyi et al., 1983) while radiatum astrocytes through their control of SOM inhibitory dendritic synapses in *stratum radiatum* and *stratum lacunosum-moleculare* could affect pyramidal cell burst firing, synaptic plasticity, and synaptic integration (Halasy et al., 1996, Lacaille et al., 1987, Pouille and Scanziani, 2004, Sik et al., 1995). While we uncovered differential gene expression between the two astrocyte populations, transcriptomic analyses with a more comprehensive range and the development of new molecular tools could advance the selective targeting and manipulation of these novel populations of astrocytes.

## Conclusion

Our results uncover the molecular, physiological and functional heterogeneity of two astrocyte populations in the CA1 hippocampus, and their local regulatory domains of SOM and PV inhibitory synapses. These domains are consistent when astrocytes are chemogenetically stimulated, and downstream of neuronal activity via optogenetic stimulation of PV and SOM interneurons. It would be worthwhile to explore further how the two astrocyte populations derive their specializations. Given the role of CA1 circuit, and astrocytes, in learning and memory, our results suggest that astrocytes heterogeneously contribute to hippocampal learning and memory.

## Methods

### Mice

All experiments were approved and conducted within the guidelines for maintenance and care of animals established by the Canadian Council of Animal Care and Université de Montréal. Male and female Pvalb-IRES-Cre (homozygous, Jackson Labs, JAX #017320) and SOM-IRES-Cre (heterozygous, Jackson Labs, JAX #013044) mice were used in this study.

### Viral constructs injection

Three to four-week-old mice were anaesthetized with i.p. injections of xylazine (5mg/kg) and ketamine (50mg/kg) and secured in a stereotaxic frame for bilateral injections of viral vectors to the dorsal CA1 hippocampus. For Ca^2+^ imaging experiments, mice received AAV2/5-gfaABC1D-cyto-GCaMP6f for expression of calcium indicator in astrocytes. For optogenetics and DREADD experiments, mice received AAV2/9-EF1a-DIO-hChR2(H134R)-EYFP for Cre-dependent expression of ChR2 in interneurons with AAV2/5-GFAP-hM3D(Gq)-mCherry or AAV2/5-GFAP-mCherry for expression of DREADD-mCherry or control mCherry in astrocytes. The injection co-ordinates were ±1.5mm ML, -2mm AP, -1.5mm DV relative to bregma. All viral vectors were sourced from the Canadian Neurophotonics Platform Viral Vector Core Facility (RRID:SCR_016477). For optogenetically-induced iLTD experiments, AAV2/9-EF1a-double floxed-hChR2(H134R)-EYFP-WPRE-HGHpA (addgene catalog #20298) was injected instead of the previously detailed Channelrhodopsin construct, in combination with AAV2/5-GFAP-mCherry. All viral vector injections were delivered between 1×10^11^ and 5×10^12^gc/ml.

### Slice preparation

Transverse hippocampal slices were collected from Pvalb-IRES-Cre and SOM-IRES-Cre mice 3-4 weeks after injections. Mice were anesthetized with isoflurane and the brain was rapidly removed and placed in ice-cold cutting solution saturated with 95% O_2_ and 5% CO_2_. The cutting solution consisted of (in mM) 120 choline chloride, 3 KCl, 1.25 NaH_2_PO_4_, 26 NaHCO_3_, 8 MgCl_2_, 17 glucose (pH 7.4 and 295-300 mOsmol). A vibratome (VT1000S, Leica) was used to obtain 300µm-thick slices, which were transferred to artificial cerebrospinal fluid (ACSF), composed of (in mM) 124 NaCl, 2.5 KCl, 1.25 NaH_2_PO_4_, 26 NaHCO_3_, 10 glucose, 1.3 MgCl_2_, 2.5 CaCl_2_ (pH 7.3–7.4, and 300 mOsmol), at 32°C for 30 minutes and maintained at room temperature for at least 30 minutes prior to experiments. For experiments examining astrocyte biophysical properties, astrocytes were labeled with the red fluorescent dye sulforhodamine 101 (SR101, 250nM, Sigma) by incubating slices in ACSF with SR101 for 30 minutes at 32°C, 15 minutes after slicing.

### Cell visualization and astrocyte calcium imaging

CA1 hippocampal neurons and astrocytes were identified using an infrared camera (LSC-70, Dage-MTI) mounted on an AXIO Examiner Z1 microscope (Zeiss) of a confocal laser scanning microscope (LSM 880, Zeiss). The microscope was equipped with a long-distance water immersion objective (20x/1.0 n.a.; W Plan-Apochromat 421452-9800, Zeiss). CA1 pyramidal cells were visually identified based on soma location and triangular morphology. Astrocytes in SP and SR were identified by their morphology, fluorescence (SR101, GCaMP6f, or mCherry), and electrical properties. eYFP expression was visualized with 488nm laser within the detection range of 499-546nm and mCherry expression was visualized with the 594nm laser within the detection range of 585-733nm. Intracellular Ca^2+^ activity of GCaMP6f-expressing CA1 astrocytes was recorded using Zen 2.1 software (Zeiss). GCaMP was excited with the 488nm laser (attenuated to 8% maximum power) and detected with 499-546nm band pass filter. Images were captured at 512*512 pixels at 1Hz for a duration of five minutes in ACSF. Astrocyte calcium signals in the processes and soma were analyzed with the astrocyte event-based software AQuA (Wang et al., 2019).

### Whole cell recordings

Whole-cell voltage clamp recordings of optogenetically-evoked IPSCs in neurons were performed using borosilicate glass pipettes (3-5 MΩ) filled with a solution containing (in mM) 130 CsCl, 10 NaCl, 10 HEPES, 1 EGTA, 0.1 CaCl_2_, 4 MgATP, 0.4 NaGTP, 10 Na_2_-phosphocreatine and 5 QX314 (pH 7.3, 290-300 mOsm). For optogenetically-induced iLTD experiments, whole-cell recordings from pyramidal neurons were obtained with an internal solution containing (in mM) 130 CsMeSO_4_, 4 NaCl, 10 HEPES, 0.5 EGTA, 10 TEA-Cl, 1 Qx314-Cl, 2 MgATP, 0.5 NaGTP (pH 7.3, 290-296 mOsm). Neurons were voltage-clamped at - 55mV for eIPSC experiments, and at 0mV for iLTD experiments (except for TBS/control experiments at -60mV). NMDA receptor antagonist AP-5 (20µM, Tocris) and the AMPA/kainate receptor antagonist NBQX (10µM, Tocris) were present in the ACSF for all neuronal recordings.

To chemogenetically activate astrocytes, clozapine N-oxide (CNO, 10µM, HelloBio) was perfused with the ACSF for 20 minutes. To chelate astrocyte intracellular calcium, astrocytes were patch-clamped in whole cell configuration for 20 minutes prior to interneuron stimulation using borosilicate glass pipettes (5-7 MΩ) containing (in mM) 45 KMeSO_4_, 10 HEPES, 4 MgCl_2_, 4 MgATP, 0.4 NaGTP, 10 Na_2_-phosphocreatine and 20 K_4_BAPTA (pH 7.3, 292-298 mOsm) in ACSF without NBQX and AP-5 prior to proceeding with neuronal recordings. The inverse agonist AM 251 (2µM, Tocris) was used to examine CB1R involvement in iLTD.

For astrocyte biophysical properties experiments, whole-cell recording of astrocytes were obtained using borosilicate glass pipettes (5-7 MΩ) containing (in mM) 125 KMeSO_4_, 10 HEPES, 4 MgCl_2_, 4 MgATP, 0.4 NaGTP, 10 Na_2_-phosphocreatine and 0.1 AF 488 hydrazide salt (Invitrogen, pH 7.3, 290-295 mOsm). Astrocyte morphology and syncytium were visualized after 20 minutes of whole-cell recording. Electrophysiological recordings of neurons and astrocytes were obtained with a Multiclamp 700B amplifier (Molecular Devices), filtered at 2kHz and digitized at 20kHz with a Digidata 1320A and acquired with pClamp 10.3 (Molecular Devices). A subset of iLTD experiments were also performed on a Nikon Eclipse E600FN with a Multiclamp 700B amplifier and DigiData 1550B with the same settings as above. Cell input resistance was monitored throughout experiments and data were excluded if holding current, series or input resistance changed beyond 20% of initial values.

### Optogenetic stimulation

Channelrhodopsin-2 was activated in PV and SOM cells by a light guide positioned above the CA1 area of the slice (473 nm blue light, custom-made light-emitting diode (LED) system or through a commercial system via a fiber-coupled LED and LED driver (M470F3 and DC4104), Thor Labs), as previously (Matos et al., 2018). The measured LED power was 7mW at the tip of a 1mm diameter light guide (custom) or 0.4mm diameter light guide (CFM14L20, Thor Labs). For each pyramidal cell recording, light-evoked IPSCs were generated by stimulation of different duration (0.1–5 ms; 0.1 Hz) to determine eIPSC input–output function. For experimental recordings in DREADD and TBS experiments, light stimulation (0.067 Hz) was adjusted to yield IPSCs of 30–40% of maximal amplitude. For DREADD experiments, eIPSCs were monitored during a control baseline period and after 20 min of CNO application. For iLTD experiments, eIPSCs were acquired every 30s, with a baseline pre-TBS of 6 minutes, and post-TBS of 30 minutes. Paired-pulse optogenetic stimulation protocol was delivered at pre-baseline and post-TBS time points, with an inter-stimulus interval of 100ms and light duration equal to the acquisition phase (0.1-2ms). Optogenetically-induced TBS was delivered in pulses at 100Hz within bursts delivered at 5Hz repeated five times, and this train was repeated five times at 30s intervals (Udakis et al., 2020).

### Assessing glial reactivity following virus injection

To confirm that the optogenetic and DREADD experiments were not confounded by glial reactivity following virus injections, three Pvalb-IRES-Cre mice were anesthetized with xylazine (25mg/kg) and ketamine (250mg/kg) 21 days after unilateral virus injection and transcardially perfused with 0.9% saline followed by paraformaldehyde (4% in phosphate buffered saline, PBS). Brain was removed and post-fixed in paraformaldehyde (4% in PBS) for 24 hours followed by 48 hours of cryoprotection (30% sucrose in PBS). Coronal sections (50µm thickness) of hippocampus were obtained via microtome cryosectioning. Sections were permeabilized in PBS with 0.3% Triton (20 min) followed by blocking in normal donkey serum (NDS; 10% in 0.1% Bovine serum Albumin, BSA, in PBS-Triton; 1 hour). Sections were incubated overnight at 4°C in fresh blocking solution containing primary antibodies of polyclonal rabbit anti-GFAP (1:1000, Dako Z0334) or polyclonal rabbit anti-IBA1 (1:1500, Wako 019-19741). Following three PBS washes, the sections were then incubated in 0.1% BSA PBST containing donkey anti-rabbit 647 (1:500) for two hours at room temperature followed by mounting (ProLong Gold antifade reagent with DAPI, P36935). Cell counts of astrocytes and microglia/macrophages were conducted in dorsal CA1.

### Analysis of spatial transcriptomics data

Spatial transcriptomics multiplexed error-robust fluorescence in situ hybridization (MERFISH) data were analyzed from the Allen Brain Cell (ABC) Atlas hosted on the Allen Institute AWS S3 repository using a dataset and metadata comprising 500 genes in 4 million cells across a single adult male C57BL/6J mouse brain (data version 20230830, metadata version 20241115; https://alleninstitute.github.io/abc_atlas_access/descriptions/MERFISH-C57BL6J-638850.html). The common coordinate framework (version 20231215; https://alleninstitute.github.io/abc_atlas_access/descriptions/Allen-CCF-2020.html), a 10µm voxel resolution established by serial 2P tomography images of 1675 young adult C57Bl6J mice was integrated to the dataset to facilitate parcellations at subclass and sublayer resolution. Detailed methods of MERFISH experimental procedures are provided in Yao et al. (2023).

Astrocytes of dorsal CA1 were filtered by removing brain sections that did not contain dorsal CA1 (retaining sections 35-42), selecting for transcriptionally defined astrocytes (class 30 Astro-Epen) and removing glial non-astrocytic clusters. The astrocytes included in analyses were exclusive to subclasses 318-321. CA1 was further refined by removing subiculum, and the CA1 SP layer was corrected by the spatial co-ordinates of excitatory pyramidal neurons (class 01 IT-ET Glut, subclass 016 CA1-ProS Glut). Based on the functional determination of the spatial distribution of astrocytes (border of 20 μm from the pyramidal layer), astrocytes through CA1 layers *stratum oriens* to *stratum lacunosum-moleculare* were remapped using this construct to identify Peri-SP astrocytes from the SR astrocytes population located within 20µm of SP through manual annotation. Gene expression data is transformed as log_2_(x+1), where x represents counts per million mapped reads. Genes were considered enriched if mean log_2_(x+1) > 1 in at least one population. Five components were used to compute the PCA and UMAP after visualizing components with a scree plot. GProfiler was used to generate gene ontology (GO) terms from the 44 DEGs. Significant biological process GO terms were ranged in fold change and p value, with term redundancy (FDR<0.05) using Revigo.

### Statistical analyses

Results are presented as mean ± standard error of the mean unless otherwise stated. Data were assessed for normality and variance assumptions with Shapiro-Wilk test and plots prior to statistical analyses. Differences in calcium events between the two astrocyte populations were analyzed with a Mann-Whitney test or unpaired t-test. Biophysical properties were analyzed with Welch’s t-test. Data from the optogenetic and chemogenetic experiments were analyzed with a paired t-test or Wilcoxin matched-pairs signed rank test when not normally distributed. TBS experiments were analyzed with a Repeated Measures 2-way ANOVA. Paired-t tests were used to assess the effect of virus injection on GFAP- and IBA1-positive cells and PPR. Information about statistical analyses for figure data is presented in Supplementary Table 1. Alpha value was set at 0.05 and all applicable tests were two-tailed. Analyses of CA1 astrocyte spatial transcriptomics were performed using two-tailed Wilcoxin t-tests with 0.01 alpha value and |log_2_FC|>0.5 threshold. Spearman correlation was used to assess similarity between CA1 astrocytes. Data were analyzed and visualized using GraphPad Prism software, except for spatial transcriptomics which was accessed, processed and analyzed using the abc_atlas_access Python package, scanpy toolkit and custom scripts.

### Data availability

The datasets and code that support the findings of this study are available from the corresponding authors upon reasonable request.

## Contributions

JCL and RR supervised the project. DC, AB, JCL and RR designed the research. AB, JCL and RR conceived the initial strategies combining chemogenetics with optogenetics and BAPTA.

AB performed and analyzed the original experiments and data providing calcium imaging evidence for functional heterogeneity (Figure 2). AB contributed to most of the data collection on Figure 3 and some experiments and analysis on Figure 4.

DC performed experiments and analyzed data of glial synaptic spatial specificity and heterogeneity (Figures 4, 5 & 6), cell/syncytium morphology experiments (Figure 1), and immunohistochemical experiments; performed some data analysis of calcium imaging experiments; and prepared all figures.

DC and KC processed, coded, analyzed and prepared Figure 7 from data available in the online repository.

EH performed electrophysiological experiments for parts of Figure 6 and for Supplementary Figures 4 and 5.

EA contributed to the experimental design of Figure 1.

DC, JCL and RR wrote the paper. All authors commented on the manuscript.

## Acknowledgements

This work was supported by grants from the Natural Sciences and Engineering Research Council of Canada (NSERC Discovery Group grant, RGPIN-2020-05264) and Canadian Institutes of Health Research (#PJT-173415). DC is supported by Fonds de recherche du Québec – Santé postdoctoral fellowships (#320029, #358284). The authors thank IL and GV for colony management.

---

**Supplementary Figure 1.**
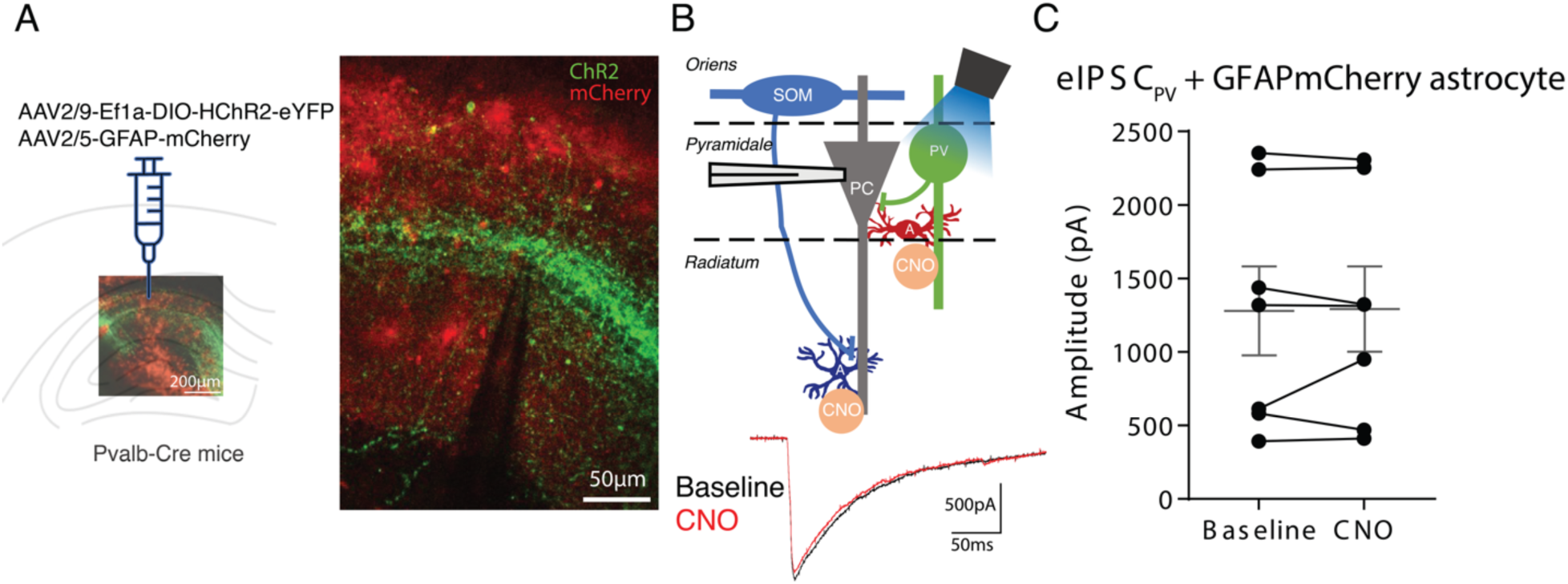
Control experiments, showing CNO (without DREADD) does not depress eIPSCs. A, Fluorescence images showing expression of GFAP-mCherry (no DREADD construct) and EF1a-DIO-hChR2-eYFP in CA1 region of Pvalb-IRES-Cre mice. B, Diagram of experimental design and representative eIPSC_PV_ in control and following CNO. C, Summary plot of eIPSC_PV_, showing no effect of CNO in these conditions (paired t-test, p>0.5, 7 cells). Dots represent individual cells, mean and SEM in grey.

**Supplementary Figure 2.**
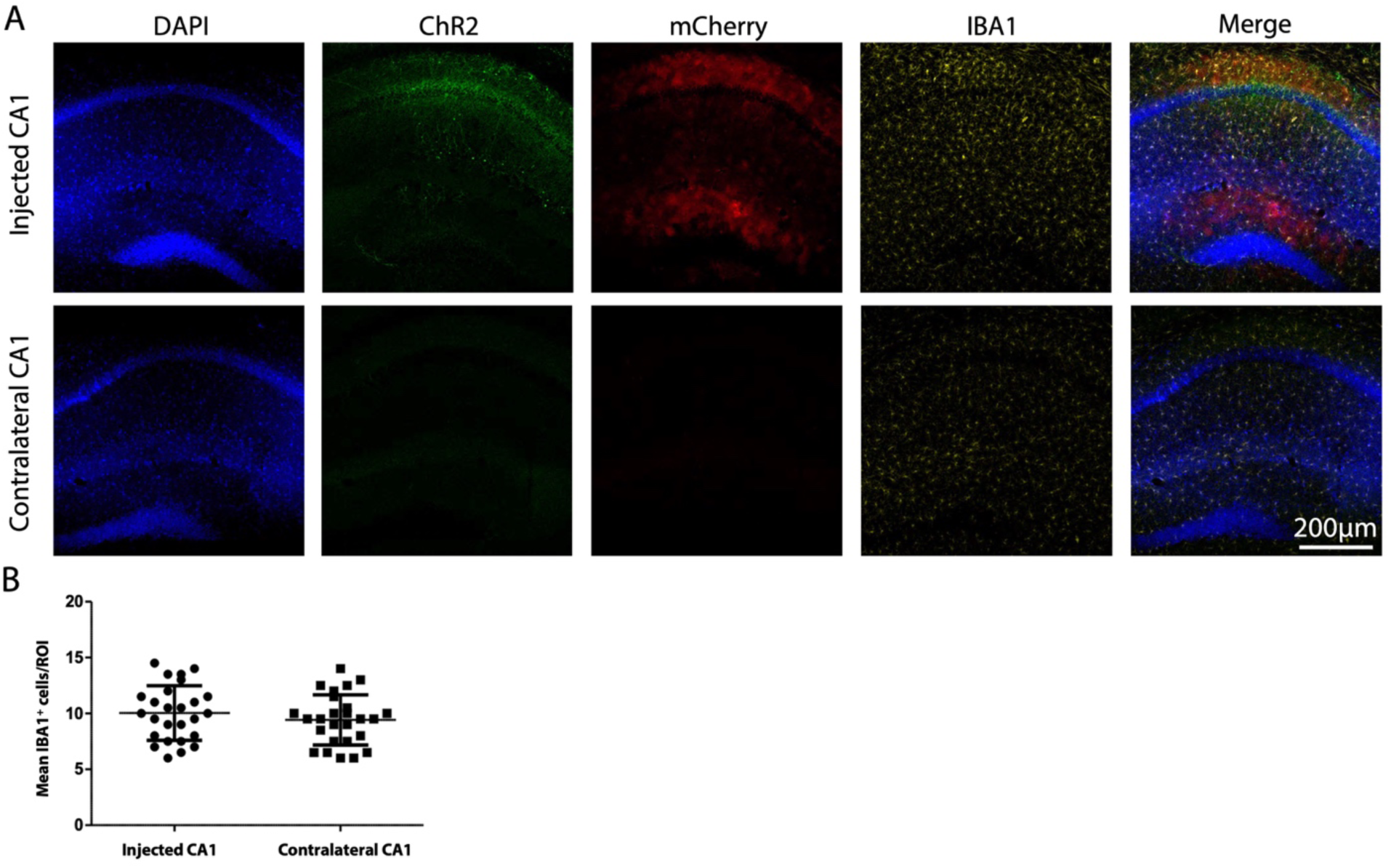
Viral construct injections do not increase the number of IBA1^+^ cells in injected CA1 hippocampus. A, Fluorescence images of immunostaining of IBA1, expression of YFP and mCherry, and DAPI, showing no immune or inflammatory response. B, Summary plot showing no difference in the number of IBA1^+^ cells in injected relative to contralateral control CA1 hippocampus (unpaired t-test, p=0.3493).

**Supplementary Figure 3.**
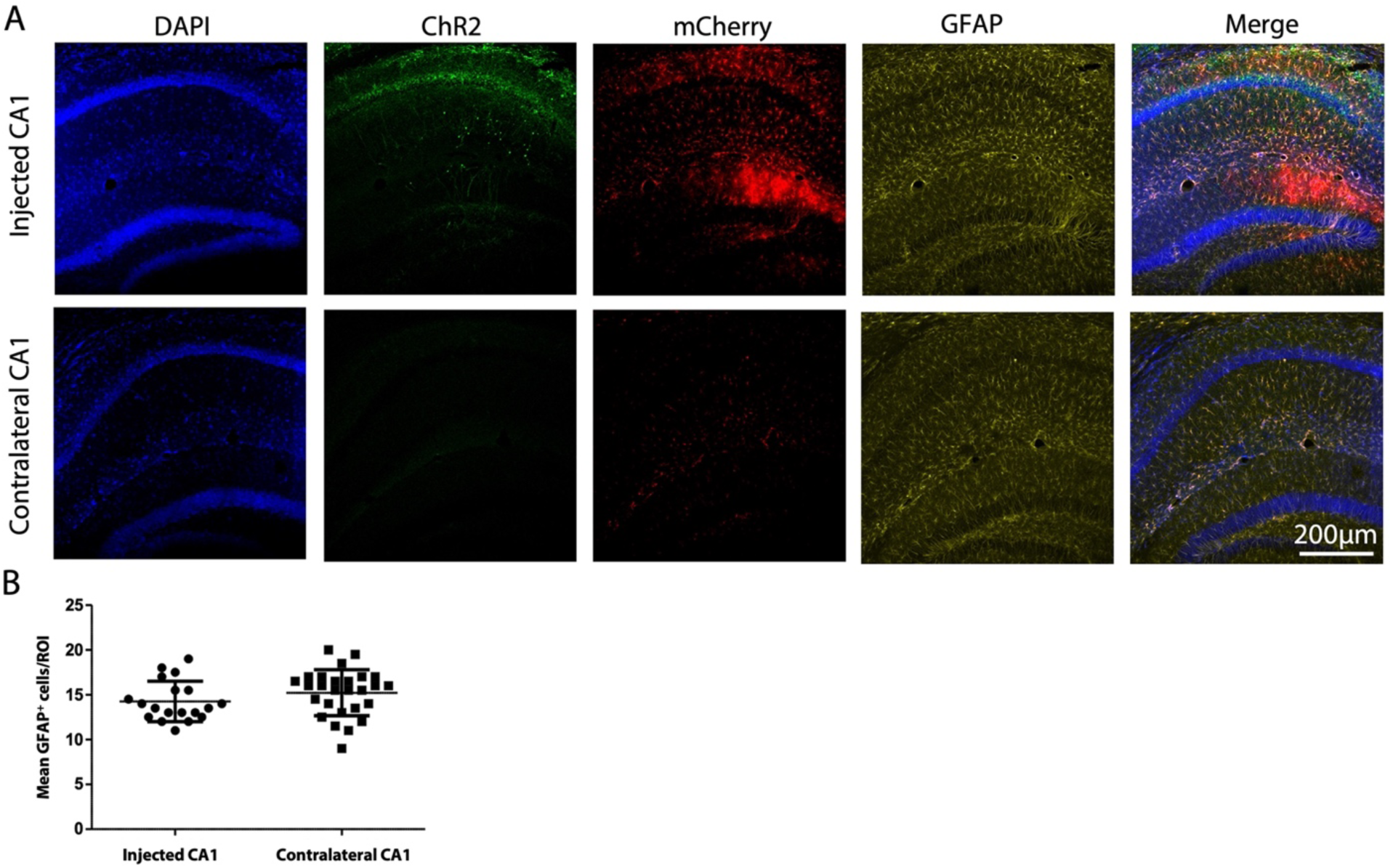
Viral construct injections do not increase the number of GFAP^+^ cells in injected CA1 hippocampus. A, Fluorescence images of immunostaining of GFAP, expression of YFP and mCherry, and DAPI, showing no glial reactivity response. B, Summary plot showing no difference in number of GFAP^+^ cells in injected relative to contralateral control CA1 hippocampus (unpaired t-test, p=0.1964).

**Supplementary Figure 4.**
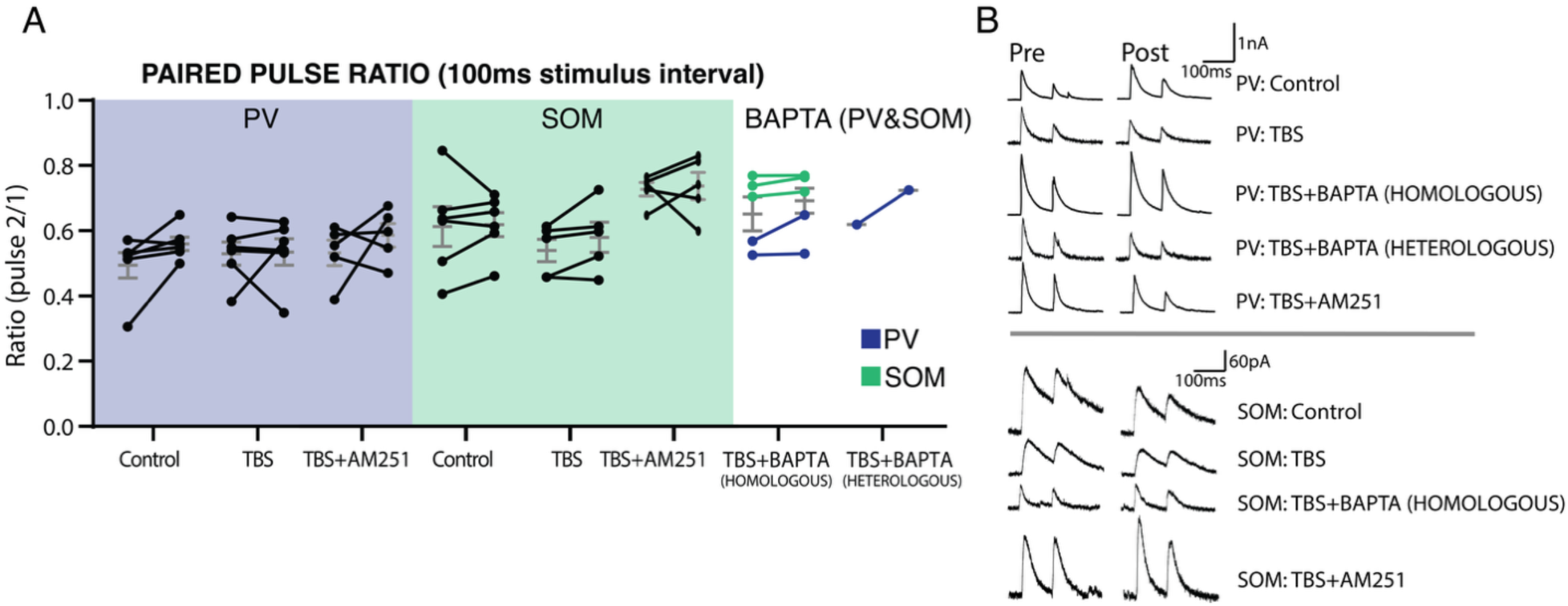
Paired-pulse ratio is not altered after TBS or BAPTA. A, Summary graph showing PPR recorded prior to baseline (left point) and 30 minutes post TBS (right point) for each group of evoked IPSCs from PV and SOM stimulation. Each line represents a recorded cell, mean and SEM provided in grey. The mean paired-pulse ratio remains unchanged with time, and after TBS, AM251, and astrocyte manipulation. Experimental numbers for each group: PV control 6 cells, PV TBS 6 cells, PV TBS+AM251 5 cells, SOM control 6 cells, SOM TBS 5 cells, SOM TBS+AM251 5 cells, TBS + Homologous BAPTA 3 SOM and 2 PV cells, PV TBS + Heterologous BAPTA 1 cell (6 cells pooled). B, Representative traces of PPR ratio for the different experimental groups.

**Supplementary Figure 5.**
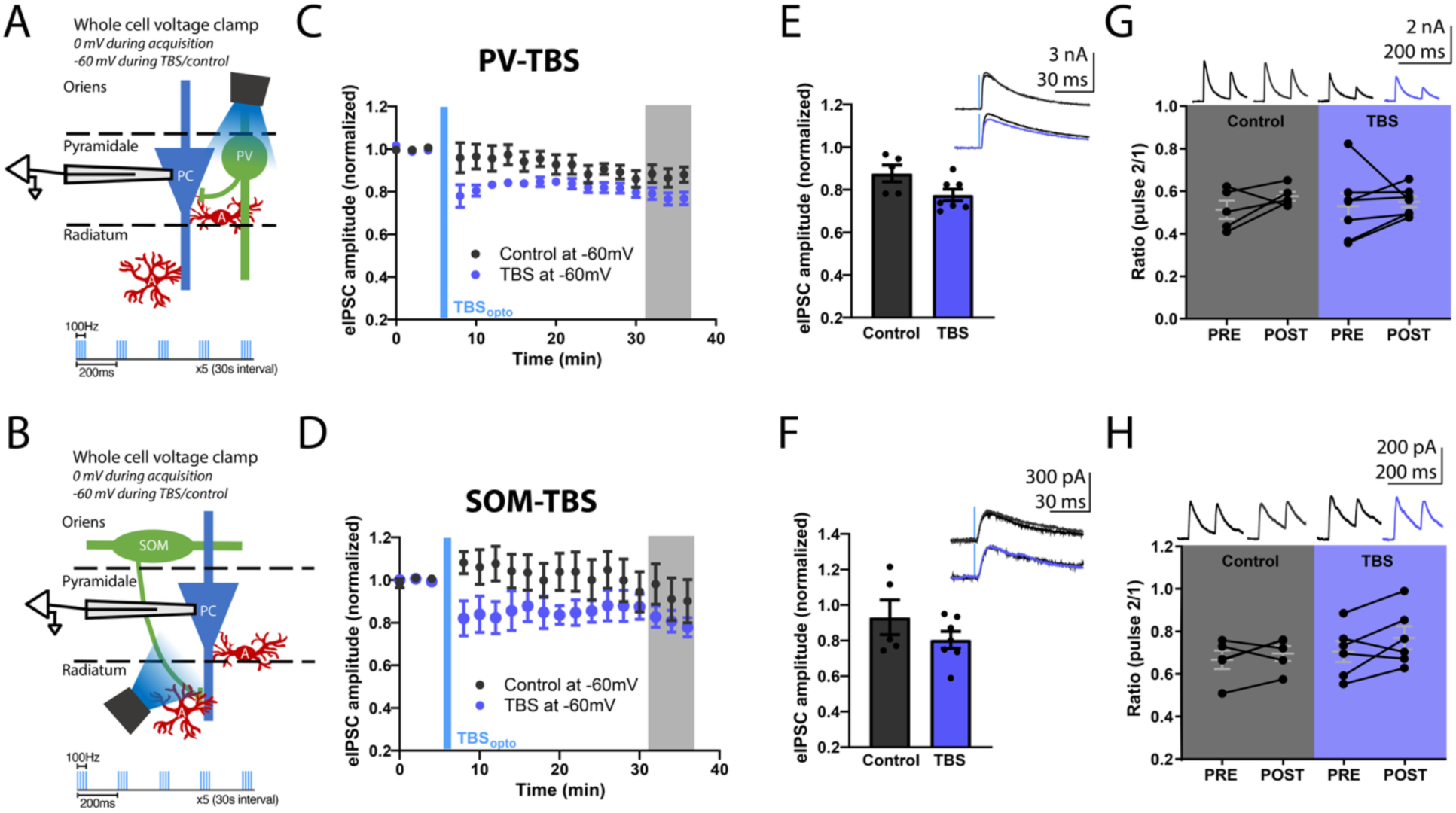
iLTD is not induced at PV or SOM synapses when pyramidal neurons are held at -60mV during TBS. A, B, Diagram of experimental design for recording eIPSC_PV_ (A) and eIPSC_SOM_ (B). C, D, Summary time plots showing normalized eIPSC_PV_ (C) and eIPSC_SOM_ (D). Data presented as mean ± SEM. E, Summary graph showing no depression eIPSC_PV_ amplitude 30 minutes after TBS when held at -60mV (Welch’s t-test p>0.05, n=5 control, 7 TBS cells). Dots represent individual cells, bars mean ± SEM. Inset shows representative traces of baseline and 30 minutes post TBS. F, Similar representation for SOM synapses. showing no depression eIPSC_SOM_ amplitude 30 minutes after TBS when held at -60mV (Welch’s t-test, p>0.05, n=5 control, 6 TBS cells). G, H, No change in PPR in these experiments with pyramidal neuron held at -60mV during TBS of PV (G) or SOM (H) interneurons. Each line represents an individual cell, mean and SEM in grey. Representative traces of PPR are provided above.

**Supplementary video 1.**
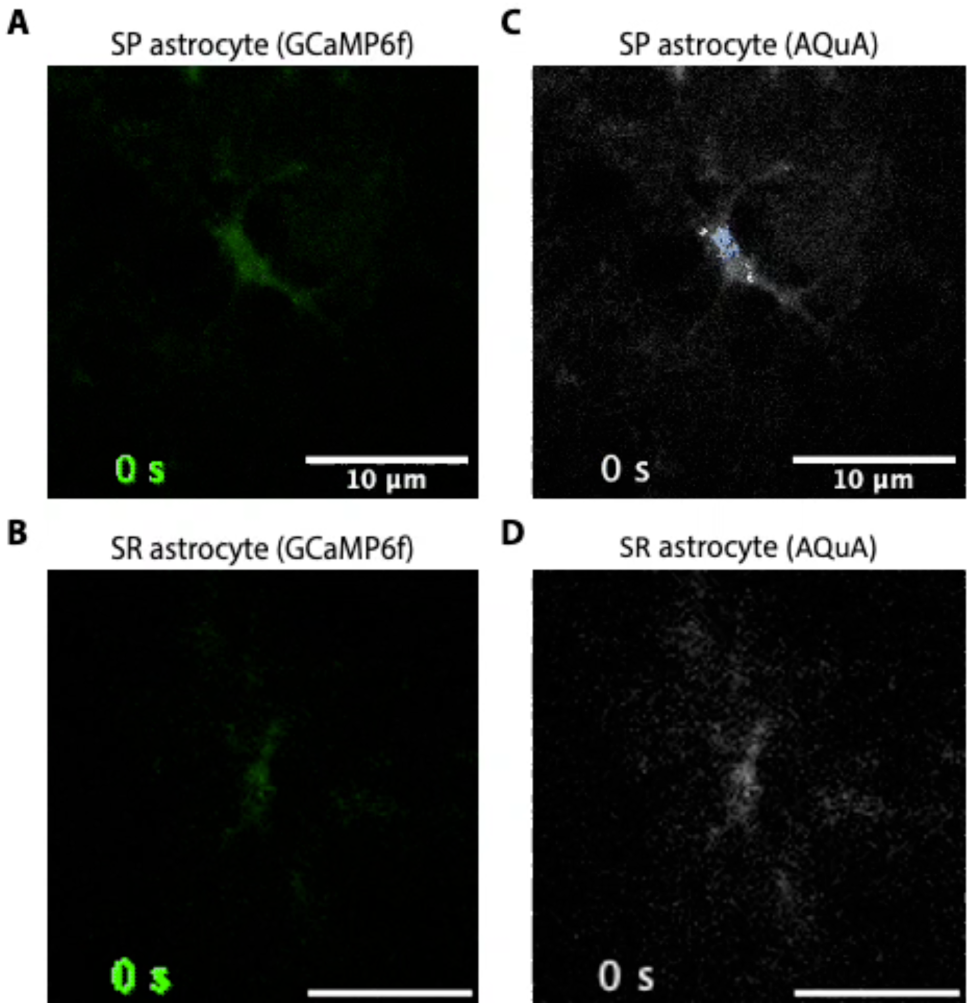
Representative movies of SP and SR astrocyte Ca^2+^ transients visualized by GCaMP6f and AQuA. A, B, Representative video of intracellular Ca^2+^ activity over 5 minutes in GCaMP6f-expressing astrocytes in SP (A) and SR (B) of CA1. C, D, AQuA movie visualizing intracellular Ca^2+^ events within the SP (C) and SR (D) astrocytes, each event is randomly pseudo-colored.

**Supplementary Table 1.**
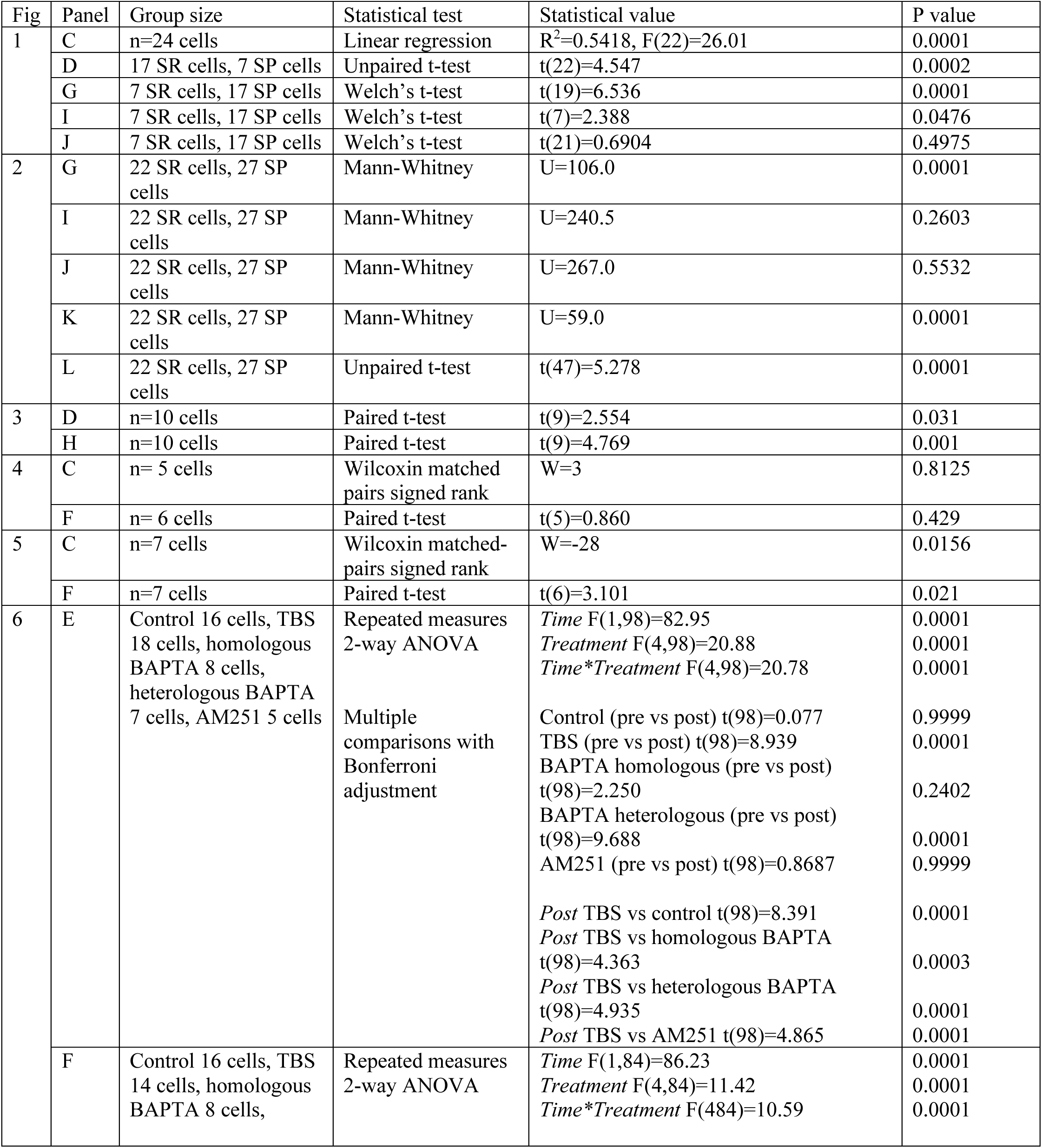
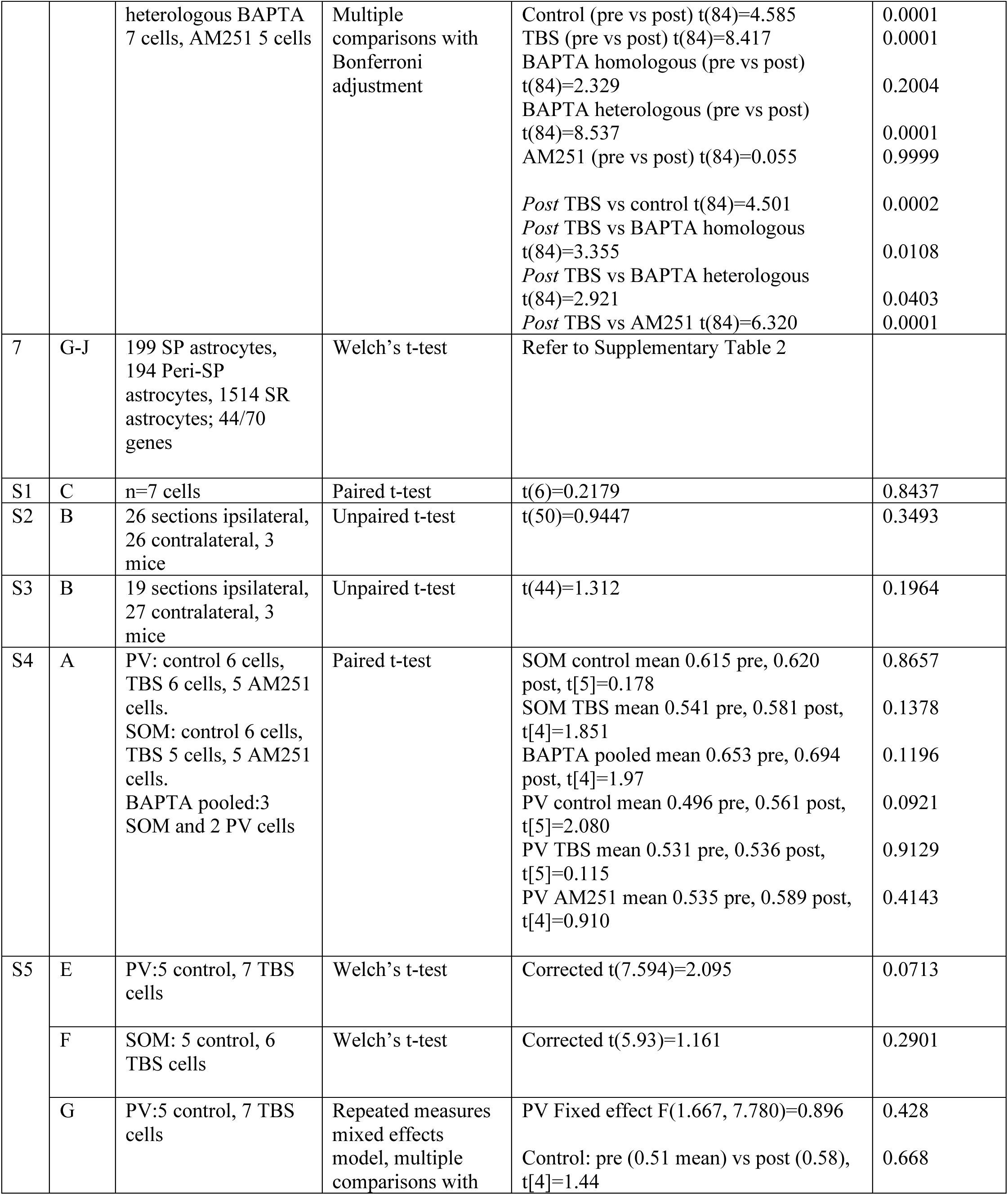
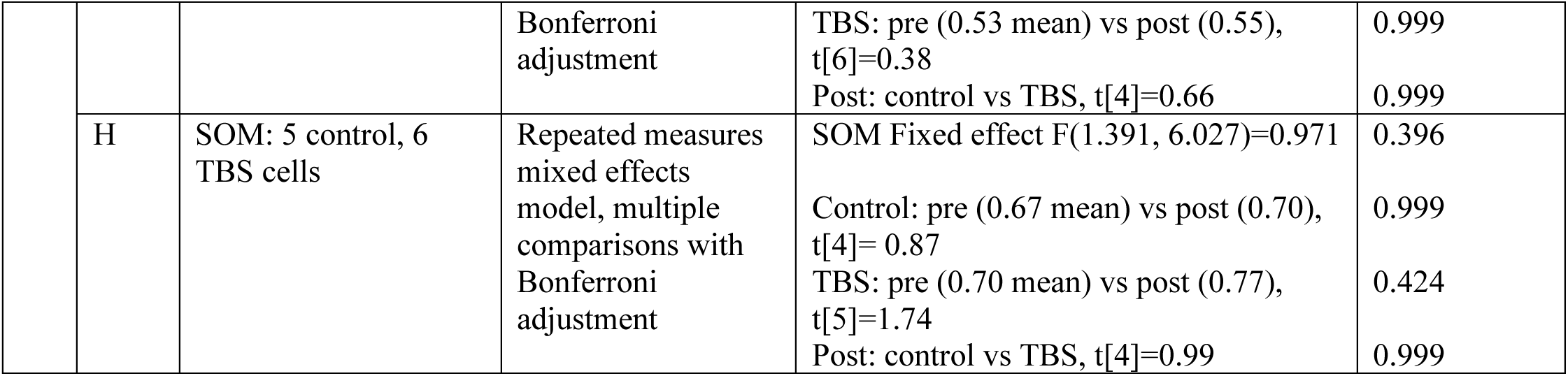
Summary of group size and statistical test information for all experiment analyses.

**Supplementary Table 2.**
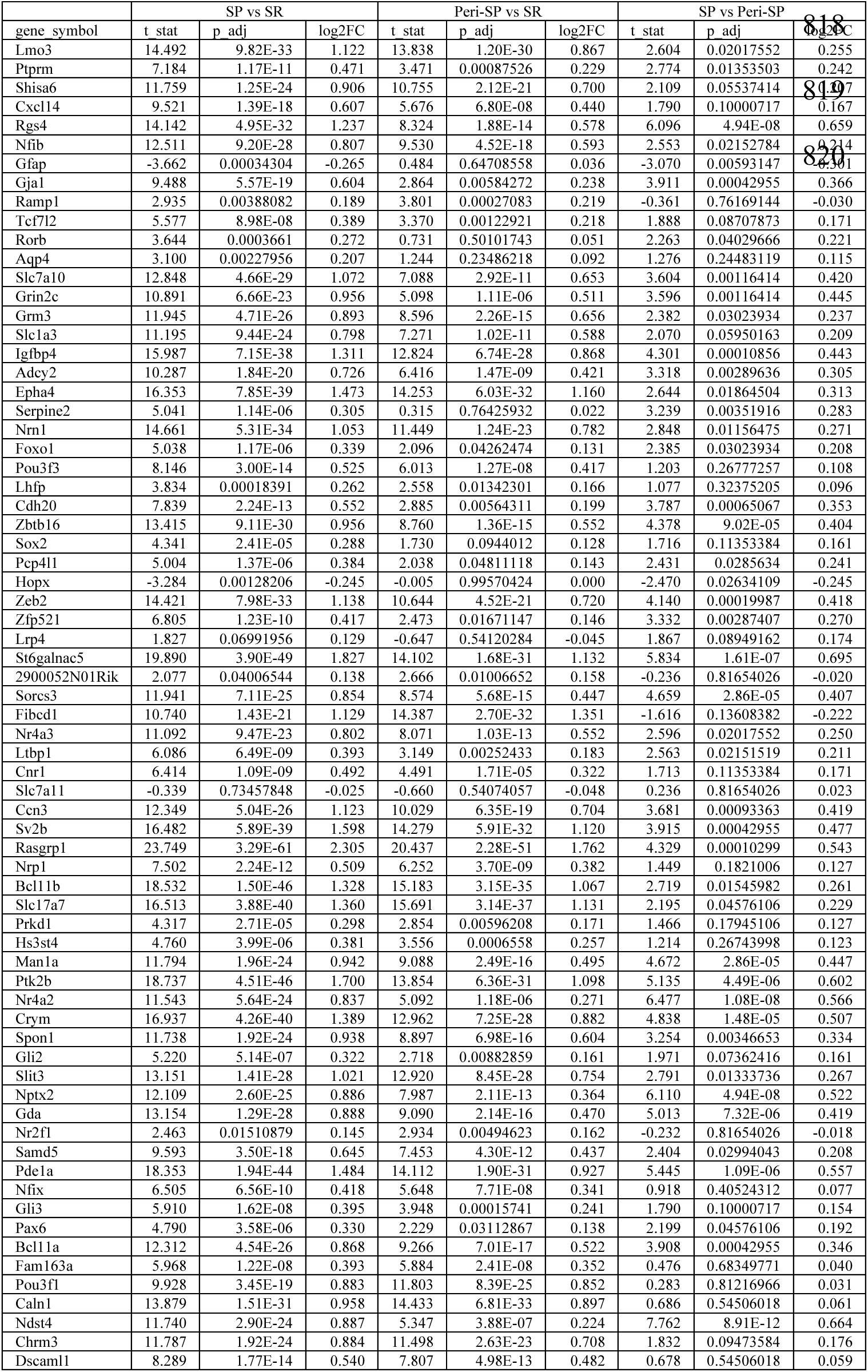
Summary of statistical values for assessment of DEG related to Figure 7.

**Supplementary Table 3.** DEG associated with top 10 GO terms for biological processes in comparison of SP and SR astrocytes.

| go term | adj_pval | # genes | gene symbols |
| --- | --- | --- | --- |
| cell morphogenesis | 3.94E-08 | 15 | GJA1, CDH20, NRP1, DSCAML1, BCL11B, NRN1, NR4A3, BCL11A, SLC1A3, ZEB2, NFIB, SLIT3, EPHA4, NR4A2, PTK2B |
| chemical synaptic transmission | 8.28E-07 | 13 | SORCS3, SHISA6, NRN1, CHRM3, SV2B, SLC1A3, GRM3, SLC7A10, EPHA4, GRIN2C, SLC17A7, PTK2B, RGS4 |
| cell-cell signaling | 1.93E-06 | 15 | SORCS3, GJA1, SHISA6, NRN1, CHRM3, SV2B, SLC1A3, GRM3, SLC7A10, CCN3, EPHA4, GRIN2C, SLC17A7, PTK2B, RGS4 |
| modulation of chemical synaptic transmission | 1.17E-05 | 11 | SORCS3, SHISA6, NRN1, SV2B, SLC1A3, GRM3, SLC7A10, EPHA4, GRIN2C, PTK2B, RGS4 |
| regulation of trans-synaptic signaling | 1.19E-05 | 11 | SORCS3, SHISA6, NRN1, SV2B, SLC1A3, GRM3, SLC7A10, EPHA4, GRIN2C, PTK2B, RGS4 |
| cell growth | 1.32E-05 | 10 | GJA1, NRP1, NRN1, BCL11A, ZEB2, SLIT3, CCN3, IGFBP4, PTK2B, RGS4 |
| nervous system development | 2.09E-05 | 18 | GJA1, POU3F3, NRP1, DSCAML1, POU3F1, BCL11B, NRN1, NR4A3, BCL11A, SLC1A3, ZEB2, NFIB, ZBTB16, SLIT3, EPHA4, NR4A2, PTK2B, RGS4 |
| regulation of signaling | 2.79E-05 | 21 | SORCS3, GJA1, NRP1, SHISA6, NRN1, CHRM3, SV2B, SLC1A3, LMO3, ZEB2, GRM3, SLC7A10, SLIT3, CCN3, EPHA4, GRIN2C, IGFBP4, NR4A2, PTK2B, RASGRP1, RGS4 |
| regulation of cell communication | 2.81E-05 | 21 | SORCS3, GJA1, NRP1, SHISA6, NRN1, CHRM3, SV2B, SLC1A3, LMO3, ZEB2, GRM3, SLC7A10, SLIT3, CCN3, EPHA4, GRIN2C, IGFBP4, NR4A2, PTK2B, RASGRP1, RGS4 |
| negative regulation of biological process | 6.21E-05 | 26 | SORCS3, GJA1, POU3F3, NRP1, DSCAML1, CXCL14, POU3F1, BCL11B, SHISA6, NR4A3, CHRM3, BCL11A, LMO3, ZEB2, NFIB, ZBTB16, SLC7A10, SLIT3, CCN3, EPHA4, GRIN2C, SPON1, NR4A2, PTK2B, RGS4, CRYM |

